# Structures reveal opening of the store-operated calcium channel Orai

**DOI:** 10.1101/284034

**Authors:** Xiaowei Hou, Shana R. Burstein, Stephen B. Long

## Abstract

The store-operated calcium (Ca^2+^) channel Orai governs Ca^2+^ influx through the plasma membrane of many non-excitable cells in metazoans. The channel opens in response to depletion of Ca^2+^ within the endoplasmic reticulum (ER). Loss- and gain-of-function mutants of Orai cause disease. Our previous work revealed the structure of Orai with a closed pore. Here, using a gain-of-function mutation that constitutively activates the channel, we present an X-ray structure of *Drosophila melanogaster* Orai in an open conformation. Well-defined electron density maps reveal that the open pore is dramatically dilated on its cytosolic side in comparison to the slender closed pore. Cations and anions bind in different regions of the open pore, informing mechanisms for ion permeation and the exquisite selectivity of the channel for Ca^2+^. Opening of the pore requires the release of cytosolic latches. Together with additional X-ray structures of an unlatched-but-closed intermediate, we propose a sequence for store-operated activation.

## Introduction

The dearth of calcium (Ca^2+^) ions in the cytosol of non-excitable metazoan cells under “resting” conditions allows transient increases in the cytosolic calcium concentration to relay internal messages and enables cells to respond to external stimuli. These cytosolic calcium signals regulate a plethora of processes including gene transcription, cell motility, exocytosis, and cell metabolism (Clapham, 2007). Short-lived increases in cytosolic [Ca^2+^] can be generated by the release of Ca^2+^ into the cytosol from the endoplasmic reticulum (ER) through ion channels such as the IP3R channel (Clapham, 2007). A second, more long-lasting elevation in cytosolic calcium occurs by the opening of Orai channels in the plasma membrane that allow Ca^2+^ ions to flow into the cell (Hogan, Lewis, & Rao, 2010). The driving force for Ca^2+^ entry is substantial: the negative voltage inside the cell is an attractive force and in the same direction as the chemical gradient for Ca^2+^ – approximately 2mM [Ca^2+^] outside the cell and 20,000–fold lower inside the cell (∼ 100 nM). Despite these driving forces Orai conducts ions approximately 1000-times slower than most channels (about 10^4^ ions per second), and this probably serves an important physiological role: it prevents overwhelming the cell with Ca^2+^ when the channel opens (Hou, Pedi, Diver, & Long, 2012; Prakriya & Lewis, 2006). How the open pore of Orai throttles the flow of Ca^2+^ is not clear. The channel is one of the most highly selective channels for Ca^2+^ but the mechanism for Ca^2+^ selectivity is not fully understood, partly because the conformation of the open pore of Orai is not known.

Calcium influx through Orai is necessary for activation of immune response genes in T cells and a range of other physiological processes (Feske, Prakriya, Rao, & Lewis, 2005; Lacruz & Feske, 2015; Prakriya & Lewis, 2015). Mutations in Orai have been implicated in a spectrum of maladies. Generally, loss-of-function mutations cause immune system dysfunction, including severe combined immunodeficiency-like disorders (Feske et al., 2006; Lacruz & Feske, 2015). Gain-of-function mutations of Orai have also been identified. These result in constitutive channel activation and have been associated with tubular aggregate myopathy and Stormorken syndromes (Lacruz & Feske, 2015).

There are three Orai proteins in humans (Orai1-3). *Drosophila melanogaster* contains one ortholog (Orai), which shares 73% sequence identity to human Orai1, and is the most studied non-human Orai channel. The channels have broad tissue distribution and are tightly regulated (Hogan et al., 2010). In the quiescent state before activation, the ion pore of Orai is closed to prevent aberrant Ca^2+^ flux through the plasma membrane. The channel is activated by the depletion of Ca^2+^ from the endoplasmic reticulum (ER), and as such it was characterized as the calcium release-activated calcium (CRAC) channel responsible for store-operated calcium entry (SOCE) before the molecular components were known (Hoth & Penner, 1992). Orai was identified as the protein that forms the channel’s pore and STIM was identified as its regulator (Feske et al., 2006; Liou et al., 2005; Prakriya et al., 2006; Roos et al., 2005; M. Vig et al., 2006; Yeromin et al., 2006; Shenyuan L Zhang et al., 2006; S. L. Zhang et al., 2005). Recent studies have uncovered the general mechanism of channel activation, which is distinct from the activation mechanisms known for other channels (reviewed in (Hogan & Rao, 2015; Prakriya & Lewis, 2015)). As a result of depletion of Ca^2+^ within the ER, STIM, which is a single-pass membrane protein resident to the ER, localizes to regions where the ER and plasma membranes are separated by only 10-20 nM. Here STIM physically interacts with cytosolic regions of Orai to open its pore. We previously determined the X-ray structure of *Drosophila melanogaster* Orai in a “quiescent” conformation with a closed ion pore but the conformational changes leading to opening and the conformation of the opened pore are unknown (Hou et al., 2012).

The X-ray structure of the quiescent conformation of Orai provides a foundation to understand the molecular basis for function (Hou et al., 2012). The channel is formed from an assembly of six Orai subunits that surround a single ion pore, which is perpendicular to the plasma membrane in a cellular setting (Figure 1A) (Hou et al., 2012). Although the oligomeric state revealed by the X-ray structure was a surprise, further studies have shown that the functional state of human Orai1 is also as a hexamer of subunits (Cai et al., 2016; Yen, Lokteva, & Lewis, 2016). Each Orai subunit contains four transmembrane helices (M1-M4) and a cytosolic M4-ext helix (Figure 1). Amino acid side chains on the six M1 helices, one from each subunit, form the walls of the pore (Figure 1B). In contrast to many ion channels, amino acid side chains establish the dimensions and chemical environment along the entirety of the pore. The M2 and M3 helices form a shell surrounding the M1 helices and shield them from the membrane. The M4 helices are located at the periphery and contain two segments, M4a and M4b, delineated by a bend at a conserved proline residue (Pro288). Following M4b, M4-ext helices extend into cytosol. The M4-ext helices from neighboring subunits interact with one another through pairwise helical coiled-coil packing, which creates a belt-like arrangement surrounding the channel on its intracellular side (Figure 1A). Mutation of the hydrophobic residues that mediate the coiled-coils has been shown to prevent channel activation by STIM, possibly by reducing the affinity for STIM (Muik et al., 2008; Navarro-Borelly et al., 2008). Because STIM also contains regions that are predicted to participate in coiled-coil helical packing, it has been proposed that STIM interacts with the M4-ext helices through such an interaction and that the structure represents a quiescent conformation prior to the binding of STIM(Hou et al., 2012). How the pore opens is not yet clear.

**Figure 1.**
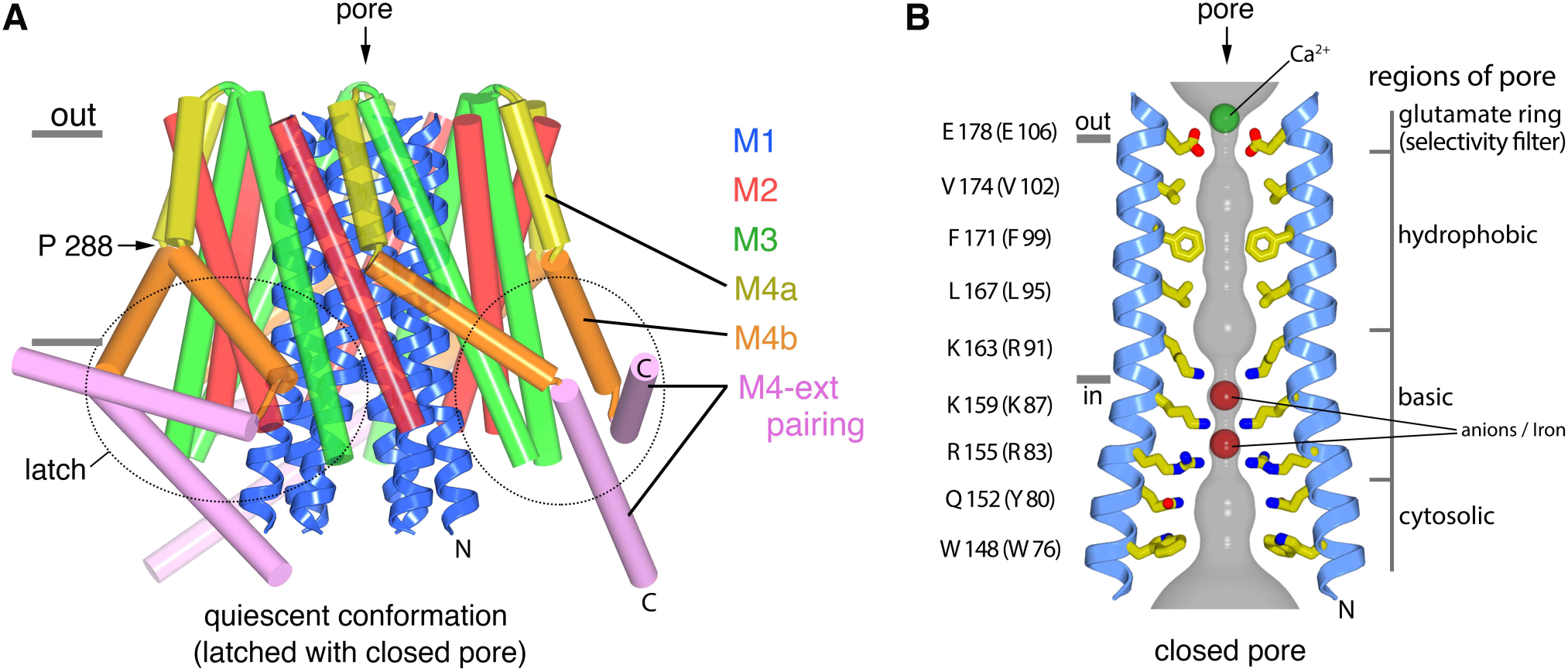
The quiescent conformation –latched with a closed pore. **a**, Overall structure of Orai, from PDB ID 4HKR, in a “quiescent” conformation(Hou, Pedi, Diver, & Long, 2012). The pore is closed; M4-ext helices pair with one another. The M1 helices are depicted as blue ribbons; other helices are cylinders. The approximate boundaries of the plasma membrane are shown as gray bars. Regions of the channel referred to as “latches” in this study are indicated as dashed ovals; the latches are comprised of interactions between M4b and M3 and the pairing of M4-ext helices. **b**, Close-up view of the closed pore. Two M1 helices are drawn as ribbons (four M1 helices are omitted for clarity). The pore is a depicted as a gray surface indicating the minimal radial distance to the nearest van der Waals contact. Amino acid side chains that form the walls of the pore are drawn as sticks and colored (yellow for carbon, blue for nitrogen, and red for oxygen). Amino acid numbering is shown for *Drosophila melanogaster* Orai without parentheses and for human Orai in parentheses. Sections of the pore are indicated. Horizontal gray bars correspond to the approximate boundaries of the membrane, although the M1 helices are shielded from the membrane by M2 and M3. A Ca^2+^ ion is indicated. Red spheres mark the location of anomalous difference electron density attributed to iron, perhaps bound as (FeCl_6_)^3-^ (Hou et al., 2012). The complex anion (IrCl_6_)^3-^ also binds in these sites(Hou et al., 2012).

The closed pore is approximately 55 Å long, narrow, and impervious to ions (Figure 1B). It contains four sections: a glutamate ring on the extracellular side that forms the selectivity filter (comprised of Glu178 residues from the six subunits), a ∼15 Å-long hydrophobic section, a ∼15 Å-long basic section, and cytosolic section (Figure 1B). Mutation of the corresponding glutamate in human Orai1 to aspartate (E106D) disrupts Ca^2+^-selectivity (Prakriya et al., 2006; Monika Vig et al., 2006; Yeromin et al., 2006). The walls of the basic section are formed by three amino acids from each of the six M1 helices that are conserved as lysine or arginine in Orai channels (Figure 1B). Mutation of one of the basic amino acids in human Orai1 (R91W, corresponding to K163W in *Drosophila* Orai) causes a severe combined immune deficiency-like disorder by preventing channel activation (Feske et al., 2006). The presence of eighteen basic residues (three from each of the six subunits) within the pore of a cation channel is highly unusual and would presumably establish an electrostatic barrier that opposes Ca^2+^ permeation in the closed state. We found that the basic region is a binding site for anion(s) that stabilize the arginine/lysine residues in close proximity (Hou et al., 2012). In the structure, an iron complex that co-purifies with the channel, which may represent (FeCl_6_)^3-^, binds at the center of the basic residues like a plug. While the physiological ligand may or may not be an iron complex, it would seem that the plug would need to be displaced to allow Ca^2+^ permeation through the pore when the channel is open.

A structure of Orai with an open pore would markedly advance our understandings of the channel but structural studies of the complex between Orai and STIM are complicated by the low affinity of the interaction and because the stoichiometry between Orai and STIM is not fully established. Gain-of-function mutations of Orai provide a potential experimental advantage for capturing a structure of an open pore because STIM would not necessarily be required for channel activation. Certain mutations of M1 residues that line the pore create constitutively active channels but these channels have reduced selectivity for Ca^2+^ in comparison to STIM-activated Orai, which suggests that their pores have non-native conformations (Beth A McNally, Somasundaram, Yamashita, & Prakriya, 2012; Yamashita et al., 2017; Shenyuan L Zhang et al., 2011). The H134A mutation of human Orai1, on the other hand, does not line the pore and has been shown to generate an activated channel with hallmarks of STIM-activated Orai, which include its high selectivity for Ca^2+^ (Frischauf et al., 2017). We introduced the corresponding amino acid substitution into *Drosophila* Orai (H206A), confirmed that it generates a constitutively active channel, and determined its X-ray structure. The structure reveals a dilated pore and conformational changes in cytosolic “latch” regions that must be released for the pore to open. In another set of experiments, we determined structures of the wild type channel and of a mutant that corresponds to one that causes immune deficiency in humans. These structures reveal an “intermediate” conformation that is in between the quiescent and opened conformations. We used additional experiments to address ion binding in the open pore that provide insight into Ca^2+^-selectivity and block by trivalent lanthanides. Together, the studies reveal mechanisms for pore opening and closing and give insight into the basis of store-operated Ca^2+^ entry.

## Results

### Activity of H206A Orai

Orai from *Drosophila melanogaster* (hereafter referred to as Orai) was selected for functional and structural studies on the basis of its good biochemical stability (Hou et al., 2012). Purified Orai containing the H206A mutation (H206A Orai_cryst_), which corresponds to the H163A gain-of-function mutation of human Orai1, was studied in proteoliposomes to assess channel activity (Figure 2). The H206A Orai_cryst_ construct is analogous to the one used to obtain the structure of the quiescent conformation, corresponds to the conserved region of Orai, and contains the regions necessary for activation by STIM (Li et al., 2007). Studying purified H206A Orai_cryst_ in proteoliposomes under divalent-free conditions, under which ionic currents through CRAC channels are more easily observed due to greater conductance of monovalent cations (e.g. Na^+^ or K^+^) than Ca^2+^ (Lepple-Wienhues & Cahalan, 1996; Prakriya & Lewis, 2006), we observed robust K^+^ flux through the channel. Ion flux was not observed for empty vesicles or through a channel without the H206A mutation (WT Orai_cryst_), as is expected without activation (Figure 2B). Similar to wild type CRAC channels, K^+^ flux was blocked by the addition of Gd^3+^ (Figure 2C) (Yeromin et al., 2006). K^+^ flux through H206A Orai_cryst_ was also inhibited by the addition of Mg^2+^ or Ca^2+^ (Figure 2C), which is in accord with the properties exhibited by STIM-activated channels and indicative of the channel’s selectivity for Ca^2+^ (Lepple-Wienhues & Cahalan, 1996; Prakriya & Lewis, 2006). Thus, as has been shown for the corresponding H134A mutation of human Orai1 (Frischauf et al., 2017), H206A Orai_cryst_ forms an open channel that recapitulates properties of STIM-activated Orai.

**Figure 2.**
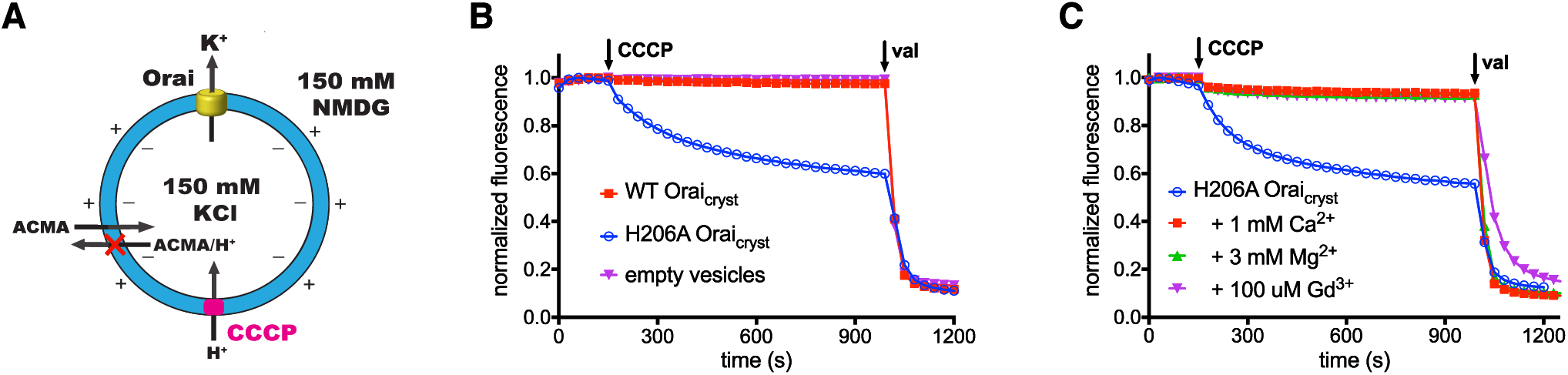
Ion flux through H206A Orai_cryst_ in liposomes. **a**, Schematic of the fluorescence-based flux assay. Vesicles containing WT or H206A Orai_cryst_ or those prepared without protein (empty vesicles) were loaded with 150 mM KCl and were diluted 50-fold into flux buffer containing a fluorescent pH indicator (ACMA) and 150 mM N-methyl-D-glucamine (NMDG) to establish a K^+^ gradient (Methods). After stabilization of the fluorescence signal (150 s), a proton ionophore (CCCP) was added. An electric potential arising from K^+^ efflux was used to drive the uptake of protons, which quenches the fluorescence of ACMA. A red “X” indicates that ACMA is not membrane-permeable in the protonated form. **b**, K^+^ flux measurements for WT and H206A Orai_cryst_. The time-dependent decrease in fluorescence observed for H206A Orai_cryst_ after the addition of CCCP is indicative of K^+^ flux. Valinomycin (val) was added after 990 s to render all vesicles permeable to K^+^ and establish a baseline fluorescence. Traces were normalized by dividing by the initial fluorescence value, which was within ±10% for each experiment. **c**, K^+^ flux through H206A Orai_cryst_ is inhibited by Ca^2+^, Mg^2+^ and Gd^3+^.

### X-ray structure of H206A Orai_cryst_ reveals an open conformation

Obtaining X-ray structural information for Orai has been challenging and capturing an open conformation of the pore especially so. Extensive optimization of crystallization conditions improved the quality of H206A Orai_cryst_ crystals from an initial diffraction limit of 20 Å resolution to 6.7 Å resolution. Despite the modest resolution of the optimized crystals, we were able to discern the conformation of the channel by calculating electron density maps using non-crystallographic symmetry averaging, which can be applied when there are multiple copies of the polypeptide in the crystallographic asymmetric unit (Bricogne, 1974). In this case, the asymmetric unit contains 24 Orai subunits, which are arranged as four complete channels. The 24-fold non-crystallographic symmetry allowed us to accurately determine the crystallographic phases and obtain electron density maps of excellent quality, which delineate all α-helices of the channel and resemble maps calculated using considerably higher resolution diffraction data (Figure 3A, B, Movie 1, Materials and Methods). All four channels in the asymmetric unit adopt the same conformation. Since side chains are not visible in the maps we collected a highly redundant dataset using an X-ray wavelength (λ=1.7085 Å) that was chosen to optimize the anomalous diffraction signal from endogenous sulfur atoms in order to locate methionine and cysteine residues within the protein (Table 1). Anomalous-difference electron-density peaks corresponding to these amino acids indicate both the validity of the atomic model and the accuracy of the crystallographic phases that were used to generate the electron density maps of the channel (Figure 3-figure supplement 1).

**Figure 3.**
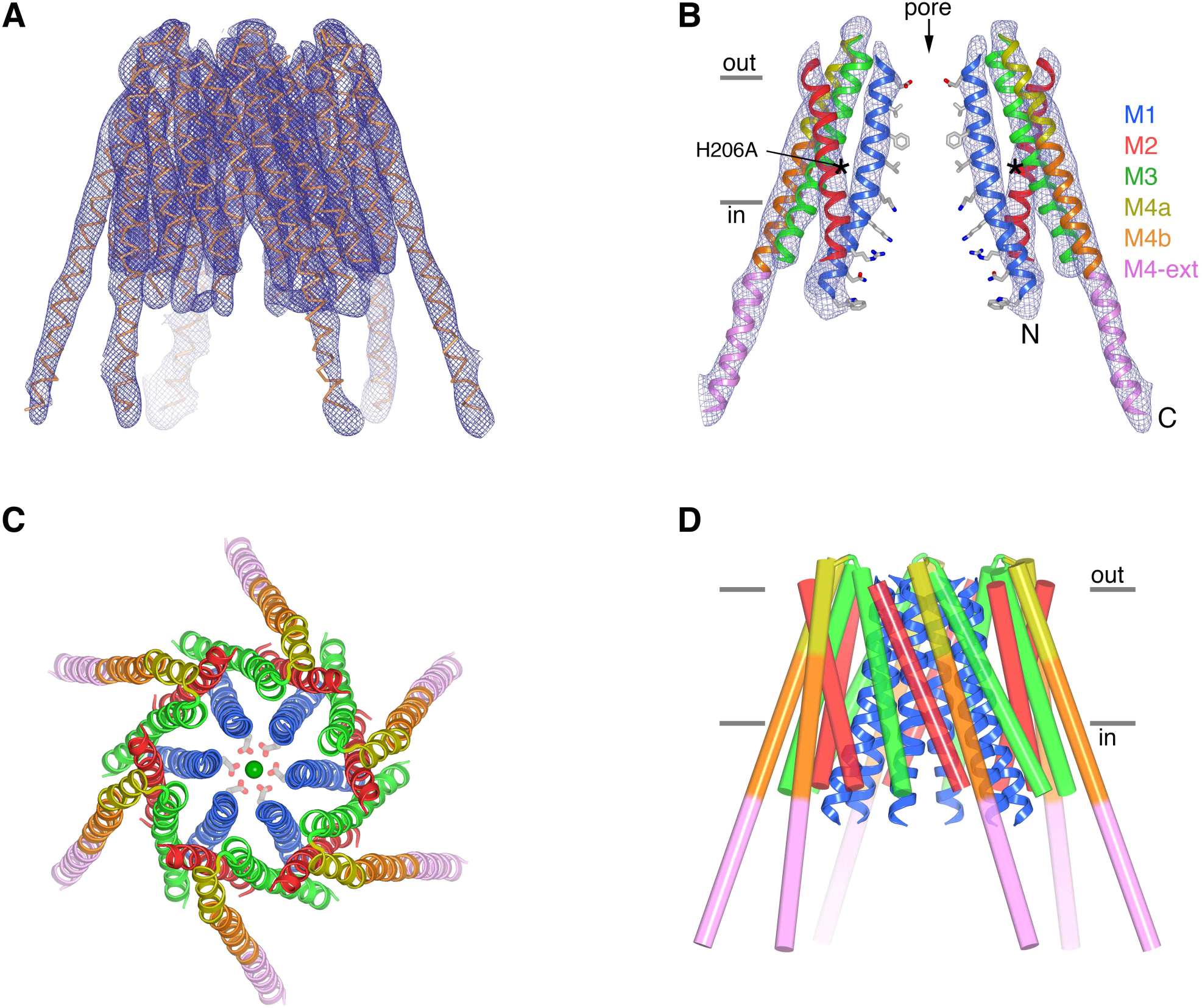
The structure of H206A Orai_cryst_ reveals an open conformation. **a**, Electron density map of H206A Orai_cryst_. The map (blue mesh, contoured at 1.4 σ, and covering one channel) was calculated from 20 – 6.7 Å using native-sharpened amplitudes and phases that were improved by 24-fold non-crystallographic symmetry (NCS) averaging, solvent flattening and histogram matching (Methods). The atomic model is shown in Cα representation. Movie 1 shows a video of this Figure. **b**, Side view showing two opposing subunits of H206A Orai_cryst_ and the same electron density map. Asterisks mark the location of the H206A substitution. Amino acid side chains on the pore are shown only for reference (sticks). Approximate boundaries of the membrane are shown as horizontal bars. Helices are depicted as ribbons and colored as indicated. **c**, Extracellular view showing the hexameric architecture. Helices are depicted as ribbons, with Glu178 side chains (sticks) and Ca^2+^ ion (green sphere) shown for reference. **d**, Overall structure, shown in the same orientation as (**a**). The M1 helices are drawn as blue ribbons and the other helices are shown as cylinders.

**Table 1.**
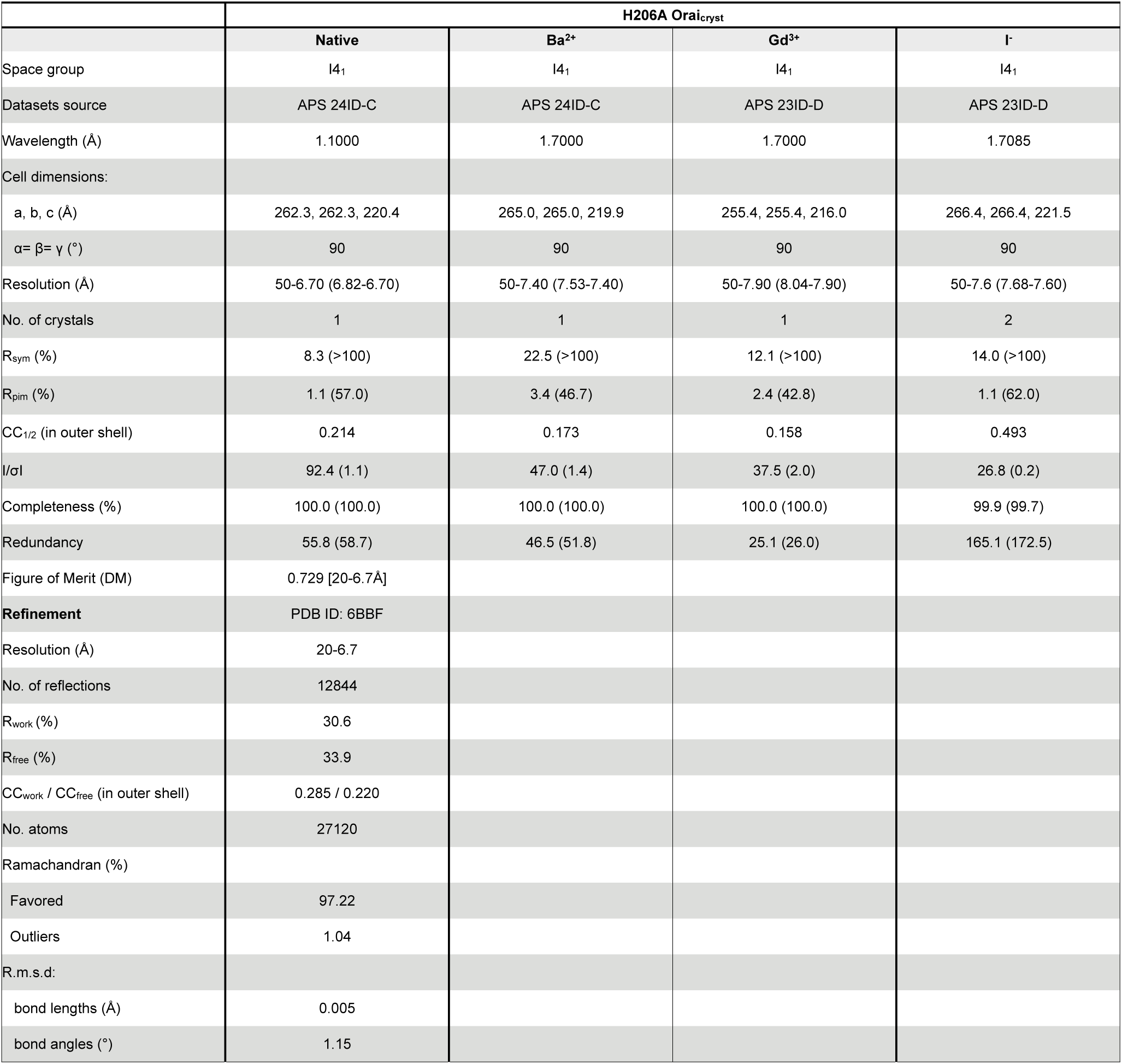
H206A Orai_cryst_ data collection, phasing and refinement statistics. Data collection statistics are from HKL3000 (Otwinowski & Minor, 1997) or XDS (I^-^ experiment) (Kabsch, 2010). R_sym_ = Σ | I_i_ - < I_i_ > | / Σ I_i_, where < I_i_ > is the average intensity of symmetry-equivalent reflections. CC_1/2_, CC_work_ and CC_free_ are defined in (Karplus & Diederichs, 2012). Phasing power = RMS (|F|/ε), where |F| is the heavy-atom structure factor amplitude and ε is the residual lack of closure error. R_cullis_ is the mean residual lack of closure error divided by the dispersive or anomalous difference. R_work_ = Σ | F_obs_ – F_calc_ | / Σ | F_obs_ |, where F_obs_ and F_calc_ are the observed and calculated structure factors, respectively. R_free_ is calculated using a subset (∼10%) of reflection data chosen randomly and omitted throughout refinement. Figure of merit is indicated after density modification and phase extension starting from 9.0 Å in DM. R.m.s.d: root mean square deviations from ideal geometry. Numbers in parentheses indicate the highest resolution shells and their statistics.

The X-ray structure of H206A Orai_cryst_ reveals an open conformation of the channel (Figure 3). The open channel is comprised of a hexameric assembly of Orai subunits surrounding a single ion pore (Figure 3). The overall architecture of the channel is similar to the quiescent conformation, with each Orai subunit containing four transmembrane helices (M1-M4). The six M1 helices, one contributed by each subunit of the channel, form the walls of the open pore. Because the secondary structure of the polypeptide surrounding the pore is α-helical, amino acid side chains on M1 establish the chemical environment along the pore. The pore is dramatically dilated on its cytosolic end, expanding by ∼ 10 Å at Lys159, in comparison to the closed pore of Orai (Figure 4) (Hou et al., 2012). The differences in between the closed and open pores taper off toward the extracellular side such that, while subtle changes may occur, the location of the M1 helix at Glu178 is indistinguishable from the closed conformation at this resolution.

**Figure 4.**
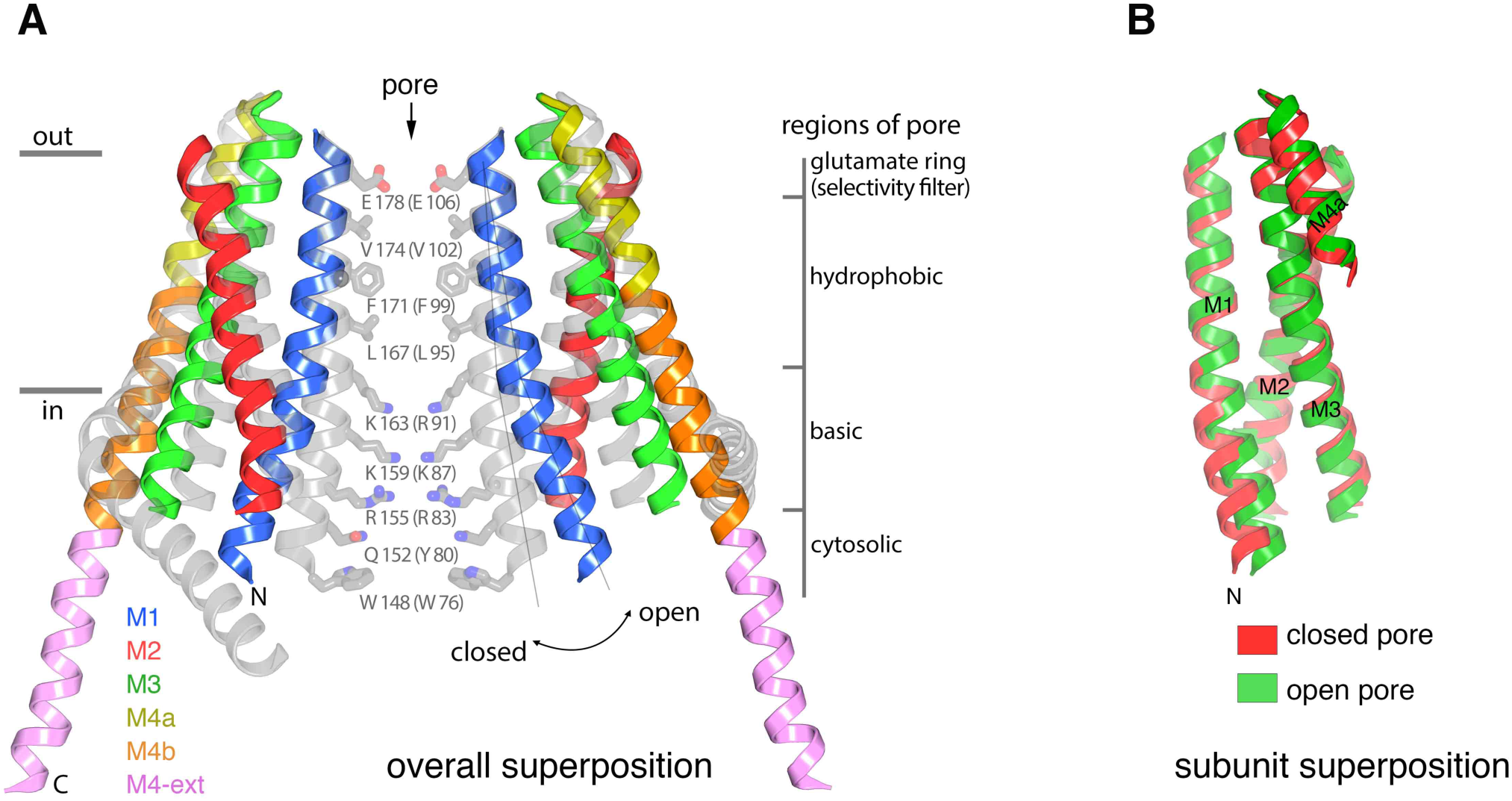
Conformational changes between the quiescent and open conformations. **a**, Superimposed structures of the quiescent (PDB ID: 4HKR) and open (H206A Orai_cryst_) conformations are drawn in ribbon representation. Two opposing subunits are shown, surrounding the pore, with the open conformation colored as indicated and the closed conformation in gray. Thin lines and a curved arrow highlight the outward rotation of subunits (with its fulcrum near Glu178) and the slight additional bend in M1. Conformational changes of M4/M4-ext are also apparent. Amino acids forming the walls of the closed pore (from the quiescent conformation) are shown as sticks, with corresponding regions of the pore indicated. Amino acids in parentheses denote human Orai1 counterparts. Horizontal bars indicate approximate boundaries of the plasma membrane. **b**, Comparison of M1-M4a from individual subunits between the quiescent and open conformations. The region of an Orai subunit spanning M1 through M4a was superimposed between the quiescent (red ribbons, PDB ID 4HKR) and open (green ribbons, H206A Orai_cryst_) conformations. The slight additional bend in M1 of the open conformation is apparent at its N-terminal end. Otherwise the M1-to-M4b region of the two subunits superimpose within the error of the coordinates of the open conformation (the root-mean-squared deviation for the Cα positions of residues 163 to 288 is 1.1 Å).

The conformational change in the pore results from an outward rigid body rotation of the M1-M4a portion of each subunit away from the central axis of the pore and a slight additional outward bend of M1 on its intracellular half (Figure 4A). The packing of M1-M4a within an individual subunit is nearly indistinguishable from the packing in the closed conformation (Figure 4B). The rigid body motion suggests that the amino acids on M1 that form the sides of the pore in the closed conformation also do so in the open conformation (Figure 3B, 4A). We cannot discern if opening involves a slight (∼20°) rotation along the helical axis of M1, which has been suggested by electrophysiological studies using cysteine mutations (Yamashita et al., 2017). On the basis of the rigid body motion from the high-resolution structure of the quiescent conformation with a closed pore, the walls of the open pore would have four sections: a glutamate ring on the extracellular side that forms the selectivity filter (comprised of Glu178 residues from the six subunits), a ∼15 Å-long hydrophobic section, a ∼15 Å-long basic section, and cytosolic section (Figure 4A).

Residue 206 is located on the M2 helix and does not line the pore (Figure 3B). In the quiescent conformation, the wild type histidine at this position forms a hydrogen bond with the side chain of Ser165, which is located on the side of M1 facing away from the pore (Figure 4-figure supplement 1A) (Hou et al., 2012). On the basis of the current structure, a histidine could be accommodated in the open conformation without steric interference, suggesting that the conformation of the pore observed for H206A Orai_cryst_ could be adopted by wild type Orai (e.g. when activated by STIM, Figure 4-figure supplement 1B). We surmise that subtle energetics involving the His206-Ser165 hydrogen bond contribute to stabilization of the closed pore and that interactions between these non-pore-lining regions of the channel influence pore opening, which is also in accord with previous studies (Frischauf et al., 2017). We postulate that the free-energy difference between the closed and open pore is on the order of a few hydrogen bonds.

### Cation binding in the open pore

To investigate potential binding sites for cations in the open pore that underlie Ca^2+^-selectivity and channel block by trivalent lanthanides, we collected X-ray diffraction data from crystals of H206A Orai_cryst_ containing Gd^3+^, which blocks the channel from the extracellular side (Aussel, Marhaba, Pelassy, & Breittmayer, 1996; Yeromin et al., 2006), and from crystals containing Ba^2+^, which is a permeant surrogate for Ca^2+^ (Hoth, 1995) that is more easily identified crystallographically. Anomalous-difference electron density maps, which pinpoint the location of these ions, contained strong density for Gd^3+^ and for Ba^2+^ in the selectivity filter (Figure 5A,B). The electron density maps could represent one or two ions that directly coordinate the side chains of the glutamate ring (Glu178 residues from the six subunits). The presence of Ba^2+^ and Gd^3+^ at this location provide evidence that the Glu178 side chains are oriented toward the pore when it is open. Ca^2+^-binding in this region likely underlies Orai’s high selectivity for Ca^2+^ under physiological conditions (e.g. extracellular [Ca^2+^] ∼2 mM). We hypothesize a selectivity mechanism that involves ion-ion interactions between two single-file Ca^2+^ ions (although the two sites may have different free energies and the electron density does not distinguish whether one or two ions are present). Block of the open channel by Gd^3+^ appears to occur by competitive binding with Ca^2+^.

**Figure 5.**
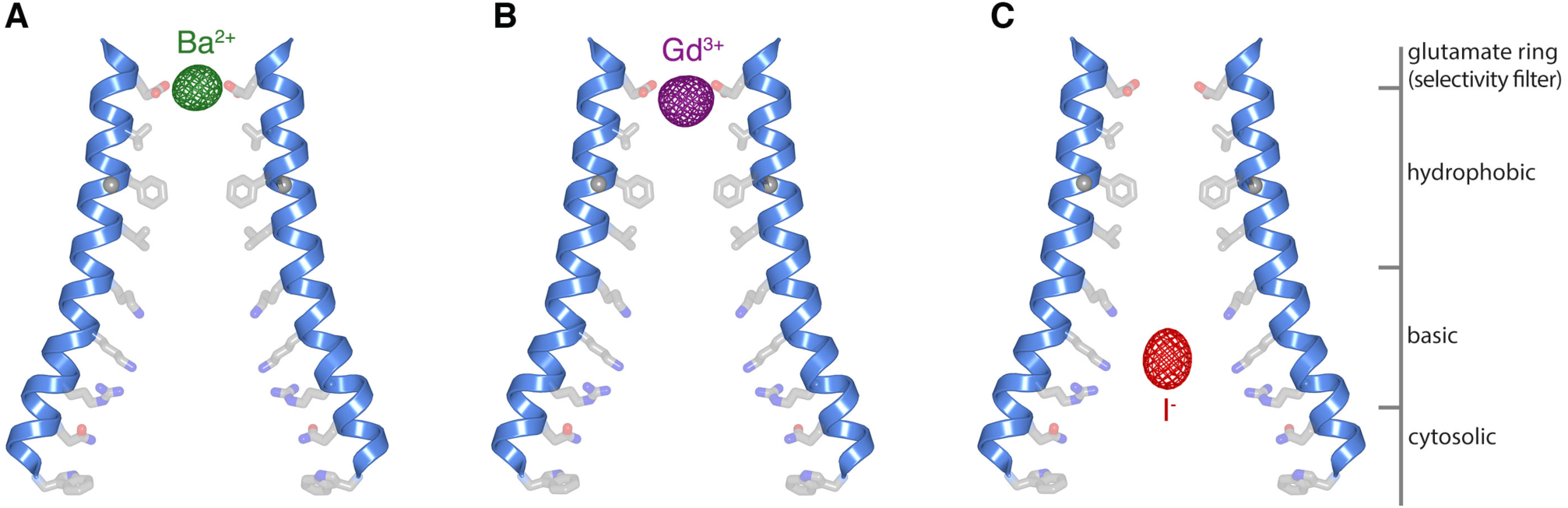
Ion binding in the open pore. **a-c**, Anomalous-difference electron density maps (mesh) for crystals of H206A Orai_cryst_ with Ba^2+^ (**a**), Gd^3+^ (**b**), and I^-^ (**c**). M1 helices of two opposing subunits are shown as ribbons. Side chains proposed to line the pore (sticks) are drawn for reference; their conformations are hypothetical. The maps are contoured at 10 σ and calculated from 25 to 9 Å for (**a-b**), and at 7 σ and calculated from 25 to 10 Å for (**c**).

We showed previously that Gd^3+^, Ba^2+^ and Ca^2+^ bind near the glutamate ring when the pore is closed (Hou et al., 2012). While the positioning of Gd^3+^ is very similar between the open and closed pores, the positioning of Ba^2+^/Ca^2+^ is noticeably different (Figure 5, Figure 5-figure supplement 1). In the closed pore, the Ba^2+^/Ca^2+^ ion binds on the extracellular side of the selectivity filter, approximately 4 Å above the ring of glutamates, whereas in the open pore, the electron density is located within the glutamate ring rather than above it (Figure 5, Figure 5-figure supplement 1). The limits of the diffraction data prevent us from discerning differences in the atomic positions of the glutamate residues between the open and closed pores but the apparent repositioning of Ba^2+^ is an indication that subtle changes occur within the selectivity filter when the pore opens. Subtle changes at the extracellular side of the pore have also been suggested by spectroscopic and electrophysiological studies when Orai is activated by STIM (Gudlur et al., 2014; Beth A McNally et al., 2012). We conclude that the transition in the pore between non-conductive and conductive conformations involves conformational changes along the length of the pore that introduce functionally important free-energy differences. These are most structurally pronounced at the cytoplasmic side but extend energetically to the selectivity filter on the extracellular side.

### Anion binding in the open pore

The basic region of the pore is highly unusual for a cation channel. We have shown previously that the basic region of the closed pore binds anions and an iron complex that co-purifies with the channel (Hou et al., 2012). Anomalous difference electron density for iron is not observed in the open pore, suggesting that the iron complex has been displaced. This is in accord with the dramatic widening of the pore in the basic region. To investigate whether anions might bind in the basic region in the open pore, and to assess if the basic amino acids contribute to the walls of the open pore, we collected diffraction data from H206A Orai_cryst_ that was crystallized in iodide (I^-^). I^-^ has similar properties to the cellularly abundant Cl^-^ anion and would be identifiable by its anomalous X-ray scattering. We observed robust anomalous difference electron density for I^-^ that is centrally located within the basic region of the open pore (Figure 5C). The presence of I^-^ there provides evidence that the basic amino acids are exposed to the pore and that anion(s) can bind in the basic region when it is open. We suspect that a few anions would coat the sides of the basic region in a cellular context. In the open conformation, the basic region is large enough to accommodate a centrally located Ca^2+^ ion that is surrounded by anions and/or water molecules. (The Cα positions of Lys^159^ residues on opposite sides of the pore are ∼24 Å apart). We hypothesize that cellular anions may shield the positive charge of the basic residues during the permeation of Ca^2+^ through the open pore.

### Mutation of the basic region

Because the basic region undergoes substantial dilation when it opens and because the eighteen basic residues (three residues from each of the six subunits) that form the walls of the pore are conserved as lysine or arginine among Orai channels, we wondered whether we could create a constitutively open channel by mutation of the residues. We substituted all three basic residues with serine (R155S, K159S, K163S) and studied the purified channel (designated SSS Orai_cryst_) in proteoliposomes using an assay to measure Na^+^ flux under divalent-free conditions (Figure 6). We chose serine because it is a small hydrophilic residue that would eliminate positive charge from this region and because the R91S mutation of human Orai1 (corresponding to K163S in Orai) forms a functional channel when expressed with STIM1 (Derler et al., 2009). We did not detect flux through SSS Orai_cryst_, suggesting that is not constitutively open. As a control, we observed Na^+^ flux through purified channels containing the V174A mutation of the hydrophobic region of the pore, which has previously been shown to produce leaky channels with diminished selectivity for Ca^2+^ (Figure 6) (Hou et al., 2012; Beth A McNally et al., 2012). We hypothesize that the modest widening of the hydrophobic region that accompanies the dramatic dilation of the basic region is critical for ion permeation through the open pore.

**Figure 6.**
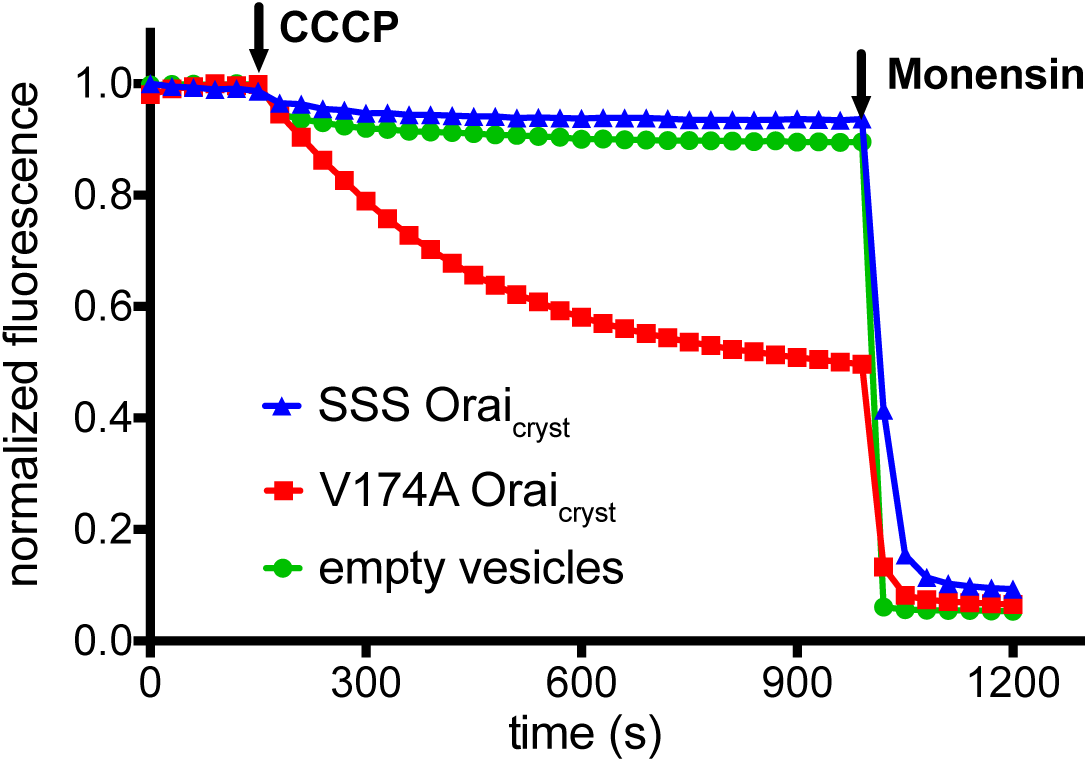
Ion flux measurements for purified channels with mutations within the hydrophobic and basic regions of the pore. “SSS Orai_cryst_” denotes the simultaneous mutation of the three basic residues to serine (R155S, K159S, and K163S). In the context of the hexameric channel, this mutant purges all eighteen basic residues from the pore. “V174A Orai_cryst_” denotes the mutation of Val174, which is located in the hydrophobic section of the pore, to alanine. Purified proteins were reconstituted into liposomes to assay for sodium (Na^+^) flux under divalent-free conditions (Methods) as described previously (Hou et al., 2012). After stabilization of the fluorescence signal (150 sec), the proton ionophore CCCP was added to the sample. A decrease in fluorescence is indicative of Na^+^ flux out of the proteoliposomes. The Na^+^ ionophore monensin was added after 990 sec to render all vesicles permeable to Na^+^ and establish the minimum baseline fluorescence. The traces were normalized to the initial fluorescence value, which was within ±10% in the experiments. Substantial fluorescence decrease is observed for V174A Orai_cryst_, indicating constitutive activity. The signal for SSS Orai_cryst_ is comparable to what is observed for liposomes without protein (“empty vesicles”).

### Conformation of M4 and M4-ext

Other differences between the quiescent and open conformations are changes in the conformations of the M4 and M4-ext helices. In the quiescent conformation, M4 and M4-ext form three helical segments: M4a and M4b, delineated by a bend in M4 at Pro288 near the midpoint of the membrane, and M4-ext, which follows a bend in a Ser306-His307-Lys308 (“SHK”) motif between M4b and M4-ext (Figure 1A). In the quiescent conformation, the M4-ext helices pair with one another through an antiparallel coiled-coil interaction (Figure 1A). In the H206A Orai_cryst_ structure, M4b and M4-ext are repositioned by straightenings of both bends such that the regions corresponding to M4a, M4b and M4-ext of each subunit form a continuous α-helix that traverses the membrane and extends ∼45 Å into the cytosolic space (Figure 3B,D and Figure 4A). The straightening of M4/M4-ext identifies the Pro288 residue and the SHK motif, both of which are conserved in Orai channels, as hinge points.

### X-ray structures of an intermediate conformation

In the crystal of H206A Orai_cryst_, the cytosolic sides of two channels face one another and the M4-ext helices of different channels interact through anti-parallel coiled-coils that are analogous to the pairing of M4-ext helices between adjacent subunits in the quiescent conformation (Figure 7A, Figure 7-figure supplement 1). To exclude the possibly that the crystal contacts in the H206A Orai_cryst_ structure were responsible for the conformational changes we observed in the pore, we determined the structures wild-type (WT) Orai_cryst_ and K163W Orai_cryst_ grown in the same crystal form (I4_1_). The K163W mutation corresponds to the R91W in human Orai1 loss of function mutation that causes a severe combined immune deficiency-like disorder (Feske et al., 2006). Well-defined electron density maps of WT and K163W Orai_cryst_ were obtained using non-crystallographic symmetry averaging of modest (6.9 and 6.1 Å, respectively) resolution diffraction data in the same manner as for the H206A Orai_cryst_ structure (Figure 8A,B). Anomalous-difference electron density for sulfur atoms of methionine and cysteine residues in WT Orai_cryst_ confirms the accuracy of the atomic model (Figure 8E). We also obtained crystals of K163W Orai_cryst_ in a P4_2_2_1_2 crystal form that diffracted X-rays to 4.35 Å resolution, which assisted with model building and was indistinguishable from the I4_1_ structures of WT and K163W Orai_cryst_ (Figure 8-figure supplement 1, and Table 2). Although the structures of WT and K163W Orai_cryst_ reveal analogous straightening of M4/M4-ext (Figure 8 and Figure 7), the pores are closed and indistinguishable from the pore in the structure of the quiescent conformation (Figure 8B,D) (Hou et al., 2012). Therefore, contacts within the crystal and the straightening of M4/M4-ext are not responsible for opening the pore. We hypothesize that additional energy would be required for the wild type pore to open, and this could be provided by the binding of STIM.

**Figure 7.**
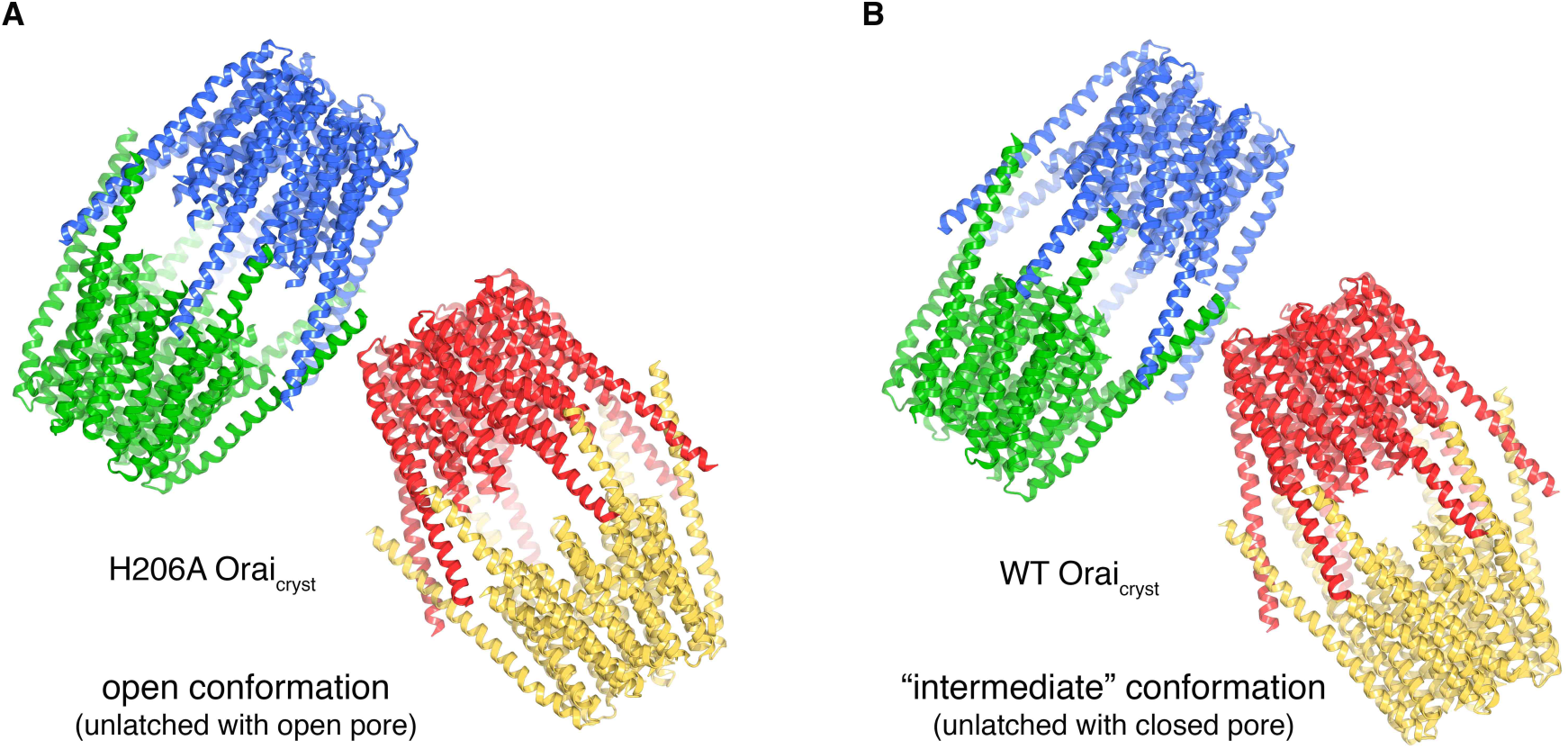
Molecular packing in the crystals of H206A Orai_cryst_ and WT Orai_cryst_. **a**, Crystal packing of H206A Orai_cryst_. The contents of the asymmetric unit, consisting of four complete channels, is shown. Each channel is colored a unique color and shown in ribbon representation. The channels interact with one another via coiled-coil interactions between their M4-ext helices. **b**, Packing of WT Orai_cryst_ in the crystal, showing the contents of the asymmetric unit, depicted analogously to **a**.

**Figure 8.**
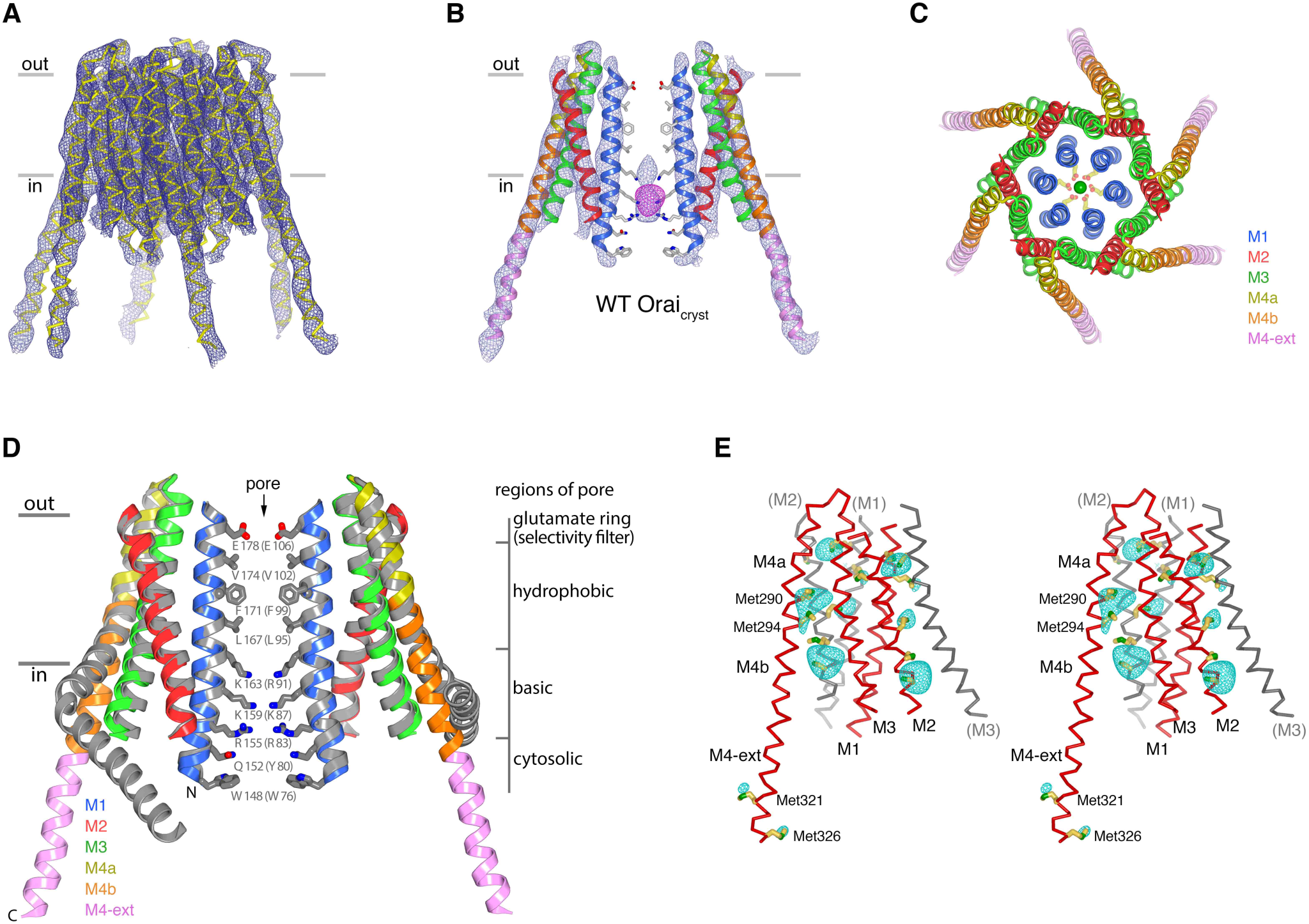
The structure of WT Orai_cryst_ reveals an intermediate conformation: unlatched with a closed pore. **a**, Electron density for WT Orai_cryst_, shown as blue mesh covering the channel (Cα representation). The map (contoured at 1.3 σ) was calculated from 20 – 6.9 Å using native sharpened amplitudes and phases that were determined by MR-SAD and improved by 24 fold NCS averaging, solvent flattening and histogram matching (Methods). **b**, Electron density, from **a** (blue mesh), covering two opposing subunits of WT Orai_cryst_ (cartoon representation, colored as indicated in **d**). Anomalous-difference electron-density (from iron) in the basic region of the pore is shown as magenta mesh (map calculated from 25 – 9 Å and contoured at 5 σ). Conformations of pore residues are based on the quiescent conformation (PDB ID 4HKR). **c**, Extracellular view of WT Orai_cryst_. Helices are drawn as ribbons and colored as indicated, with Glu178 side chains (sticks) and Ca^2+^ ion (green sphere) shown for reference. **d**, Superposition of the crystal structure of the quiescent conformation (PDB ID 4HKR) and the structure of the intermediate conformation (WT Orai_cryst_). Two subunits of each channel are shown. The quiescent conformation is gray; the structure of WT Orai_cryst_ is shown in colors. Amino acids lining the pore of the quiescent conformation are shown as sticks. A slight outward displacement of the intracellular side of M3 is observed in the structure of WT Orai_cryst_; otherwise the conformations of M1-M4a are indistinguishable within the resolution limits of the diffraction data (RMSD for Cα positions 148 to 288 is 0.9 Å). **e**, Anomalous-difference electron-density peaks at cysteine and methionine residues confirms the amino acid register of the WT Orai_cryst_ structure (stereo representation). An anomalous-difference electron-density map was calculated from 25 to 9 Å resolution from data collected with λ = 1.738 Å X-rays (Extended Data Table 2) using anomalous differences as amplitudes and phases from (**a**). This map was then averaged in real-space according to the 24-fold NCS symmetry to yield the map shown. The map is contoured at 5 σ (cyan mesh) and shown in the vicinity of a subunit of Orai (red Cα trace). Methionine and cysteine residues are shown as sticks (colored yellow for carbon and green for sulfur atoms). Methionine residues on M4b and M4-ext are labeled. Portions of neighboring Orai subunits (gray Cα traces) are shown for reference with their helices labeled in parentheses. Side chain conformations are hypothetical.

**Table 2.** Data collection, phasing and refinement statistics for WT and K163W Orai_cryst_. Data collection statistics are from HKL3000 (Otwinowski & Minor, 1997). R_sym_ = Σ | I_i_ - < I_i_ > | / Σ I_i_, where < I_i_ > is the average intensity of symmetry-equivalent reflections. CC_1/2_, CC_work_ and CC_free_ are defined in (Karplus & Diederichs, 2012). Phasing power = RMS (|F|/ε), where |F| is the heavy-atom structure factor amplitude and ε is the residual lack of closure error. R_cullis_ is the mean residual lack of closure error divided by the dispersive or anomalous difference. R_work_ = Σ | F_obs_ – F_calc_ | / Σ | F_obs_ |, where F_obs_ and F_calc_ are the observed and calculated structure factors, respectively. R_free_ is calculated using a subset (∼10%) of reflection data chosen randomly and omitted throughout refinement. Figure of merit is indicated after density modification and phase extension starting from 8.0 Å in DM. R.m.s.d: root mean square deviations from ideal geometry. Numbers in parentheses indicate the highest resolution shells and their statistics.

|  | K163W Orai<sub>cryst</sub> | K163W Orai<sub>cryst</sub> |  |  | WT Orai<sub>cryst</sub> |
| --- | --- | --- | --- | --- | --- |
|  | Native | Native | Derivative 1<br>PCMB | Derivative 2<br>PIP | Native |
| Space group | P4<sub>2</sub>2<sub>1</sub>2 | I4<sub>1</sub> | I4<sub>1</sub> | I4<sub>1</sub> | I4<sub>1</sub> |
| Datasets source | NSLS X25 | NSLS X25 | NSLS X29 | NSLS X29 | NSLS X25 |
| Wavelength (Å) | 1.1000 | 1.1000 | 1.0074 | 1.0712 | 1.738 |
| Cell dimensions: |  |  |  |  |  |
| a, b, c (Å) | 118.7, 118.7, 122.4 | 247.5, 247.5, 210.2 | 246.1, 246.1, 210.0 | 250.6, 250.6, 211.8 | 250.4, 250.4, 210.4 |
| α= β= γ (°) | 90 | 90 | 90 | 90 | 90 |
| Resolution (Å) | 60-4.35 (4.42-4.35) | 50-6.10 (6.20-6.10) | 50-6.10 (6.20-6.10) | 50-6.90 (7.02-6.90) | 50-6.9 (7.02-6.90) |
| No. of crystals | 1 | 1 | 1 | 1 | 1 |
| R<sub>sym</sub> (%) | 6.0 (>100) | 5.9 (>100) | 5.7 (>100) | 12.0 (>100) | 9.4 (>100) |
| R<sub>pim</sub> (%) | 1.7 (55.7) | 1.3 (>100) | 1.9 (>100) | 3.1 (>100) | 3.0 (>100) |
| CC<sub>1/2</sub> (in outer shell) | 0.265 | 0.376 | 0.378 | 0.170 | 0.123 |
| I/σI | 61.1 (1.0) | 69.0 (1.0) | 49.3 (0.7) | 37.0 (.07) | 39.1 (0.5) |
| Completeness (%) | 100.0 (100.0) | 100.0 (100.0) | 99.9 (100) | 99.9 (100) | 99.7 (99.8) |
| Redundancy | 16.4 (16.9) | 22.9 (23.9) | 21.5 (20.5) | 16.3 (16.7) | 11.0 (11.6) |
| **MIRAS Phasing** |  |  |  |  |  |
| No. of sites |  |  | 24 | 24 |  |
| Phasing power (iso/ano) |  |  | 0.523 / 0.559 | 0.405 / 0.686 |  |
| R<sub>cullis</sub> (iso/ano) |  |  | 0.764 / 0.945 | 0.952 / 0.919 |  |
| Figure of Merit (DM) | 0.627 [20-4.35Å] | 0.629 [20-6.1Å] |  |  | 0.777 [20-6.9Å] |
| **Refinement** | PDB ID: 6BBI | PDB ID: 6BBH |  |  | PDB ID: 6BBG |
| Resolution (Å) | 20-4.35 | 20-6.1 |  |  | 20-6.9 |
| No. of reflections | 6035 | 14682 |  |  | 10287 |
| R<sub>work</sub> (%) | 30.6 | 31.4 |  |  | 33.4 |
| R<sub>free</sub> (%) | 32.9 | 34.0 |  |  | 35.4 |
| CC<sub>work</sub> / CC<sub>free</sub> (in outer shell) | 0.487 / 0.441 | 0.346 / 0.333 |  |  | 0.332 / 0.414 |
| No. atoms | 3338 | 27360 |  |  | 27240 |
| Ramachandran (%) |  |  |  |  |  |
| Favored | 95.5 | 97.2 |  |  | 96.6 |
| Outliers | 0.24 | 1.01 |  |  | 0.93 |
| **R.m.s.d:** |  |  |  |  |  |
| bond lengths (Å) | 0.006 | 0.005 |  |  | 0.005 |
| bond angles (°) | 1.15 | 1.06 |  |  | 1.07 |

The structures of WT and K163W Orai_cryst_ resemble to be an “intermediate” because their pores are closed like the quiescent conformation and their M4/M4-ext regions are like those observed in the open conformation. In this intermediate conformation, the M1-M4a portions of the channel adopt indistinguishable conformations relative to the quiescent conformation (Figure 8D). As such, the dimensions and chemical nature of the pore appear unchanged from the quiescent conformation. As in the quiescent conformation, anomalous difference electron density for an iron complex is observed in the basic region of the pore in the structures of the intermediate conformation (Figure 8B, Figure 8-figure supplement 1C). The density is positioned roughly in the center of the basic region in the structure of WT Orai_cryst_ and located in the lower portion of it in the structures of the K163W mutant, which removes the top ring of basic residues from the pore (Figure 8B, Figure 8-figure supplement 1C). The K163W mutation has no other apparent effect on the intermediate conformation, and this is analogous to observations from structures of the quiescent conformation with and without the K163W mutation (Hou et al., 2012). The presence of the anomalous density in the basic region for both the quiescent and intermediate conformations is another indication that the pores share the same closed conformation. Because the quiescent and intermediate conformations have been obtained using nearly identical protein constructs (differing only by two amino acid substitutions in the hyper variable extracellular M3-M4 loop, Materials and Methods), we suspect that there is an equilibrium between bent and unbent conformations of M4/M4-ext. The molecular constraints of crystallization may bias the equilibrium, and STIM binding may do so in a cellular context.

### Unlatching of M4b/M4-ext is necessary for pore opening

Comparison of the structures of the quiescent, intermediate and open conformations indicates that the M4b and M4-ext regions must undergo conformational changes from the quiescent conformation for the pore to open. In the quiescent conformation, the three sets of paired M4-ext helices create an assembly surrounding the intracellular side of the channel (Figure 1A). Bends at Pro288 and in the SHK motif are necessary for this configuration. Because of the bend at P288, M4b interacts with M3 (Figure 1A, Figure 9A). In the open structure, the interaction between M4b and M3 is no longer present due to the repositioning of M4b that is enabled by unbending at Pro288 and the unpairing of M4-ext helices (Figure 9C). If the interaction between M4b and M3 of the quiescent conformation were present, or if the M4-ext helices were paired, the rigid body motion of M1-M4a that underlies pore opening could not occur due to steric interference between M3 and M4b (Figure 9D). We conclude that the paired M4-ext helices and the concomitant interactions between M4b and M3 of the quiescent conformation constitute “latches” that must be released for the pore to open. In belt-like fashion, the latches constrain the outer diameter of the intracellular portion of the channel and prevent the widening observed for the open pore. Thus, when the latches are fastened they stabilize the pore in a closed conformation. Complete straightening of the P288 and SHK bends, like what is captured in the structures of H206A, WT, and K163W Orai_cryst_ presented here, may not be necessary for the pore to open because there could be enough space for pore dilation without complete straightening. The hinges may provide flexibility to the M4b and M4-ext helices when the latches are released. We hypothesize that the straightened conformations of the M4/M4-ext helices in the crystal structures are one conformation of these mobile regions along a continuum of unlatched conformations that would permit, and necessarily precede, the opening of the pore. The structures of the intermediate conformation reveal that release of the latches does not necessarily open the pore: the pore is closed despite the M4b and M4-ext helices adopting the same conformation that they do in the open conformation. Thus, while necessary, unlatching is not sufficient to open the pore.

**Figure 9.**
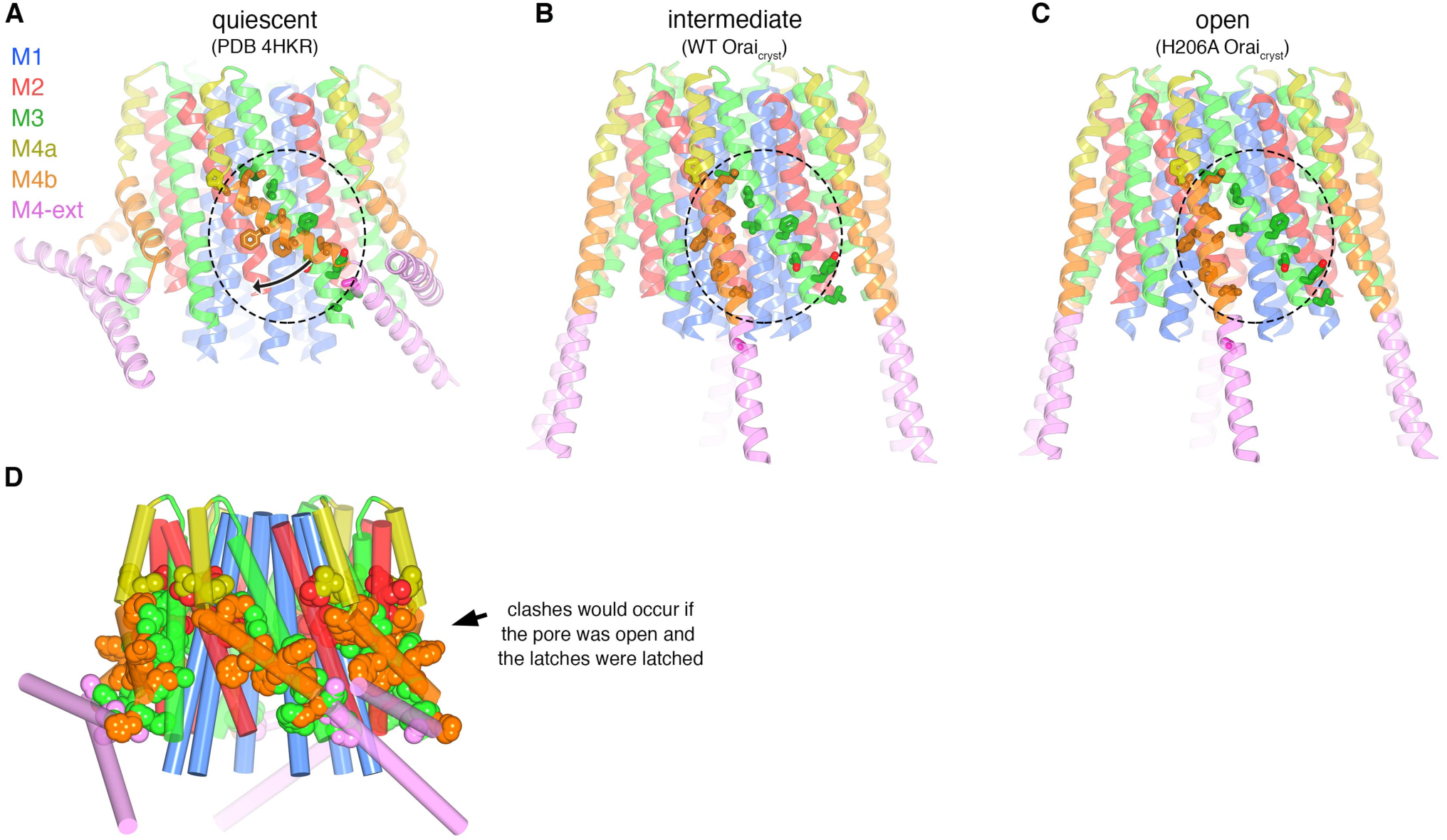
Unlatching is necessary for pore opening. **a**, Quiescent conformation (PDB ID: 4HKR) highlighting interactions between M4b and M3. The channel is shown in cartoon representation. On one of the Orai subunits, amino acids in the interface between M4b (orange) and M3 (green), which is highlighted by a dashed oval, are shown as sticks and colored accordingly. This interface exists for all six subunits and is stabilized by the pairing of M4-ext helices (pink). An arrow denotes the movement of M4b between (**a**) and (**b**). **b**, Conformation of WT Orai_cryst_ showing the released latches and closed pore. The amino acids that had been in the interface between M4b and M3 in the quiescent conformation are drawn as sticks, with the dashed region showing the same region as in (**a**). **c**, Open conformation (H206A Orai_cryst_), depicted as in (**b**). **d**, Unlatching is necessary for pore opening. A hybrid atomic model of the channel was generated using the conformations of the M1, M2 and M3 helices from the open conformation and the conformations of M4a/b and M4-ext from the quiescent conformation. In this model (shown in cartoon representation), molecular clashes exist between M4b and M3. Amino acids involved in the clashes are depicted as space-filling spheres (colored according channel region: red, M2; green, M3; yellow, M4a; orange, M4b; pink, M4-ext). These steric hindrances would prevent opening of the pore while the latches are fastened. With unlatching, the repositioning of the M4b helix would allow the outward motion of M1-M3 that opens the pore.

In a cellular context, mutations that cause release of the latches, that is, those that destabilize the quiescent conformation and/or favor straightening of the M4/M4-ext helices, may appear as activating mutations. Congruently, Pro245 of human Orai1, which corresponds to Pro288 of Orai, has been characterized as a residue that helps stabilize a closed state of the channel - mutation to any other residue, which would favor straightening of the bend at this position, has an activating phenotype (Nesin et al., 2014; Palty, Stanley, & Isacoff, 2015). Further, certain mutations of human Orai1 within and around the SHK hinge also create active channels (Zhou et al., 2016). Because unlatching is necessary, but not sufficient, to open the pore, these mutations may increase the probability that the channel is open in the absence of STIM and/or they may increase the binding affinity for STIM.

## Discussion

Studies have shown that the M4-ext region of Orai interacts with a cytosolic portion of STIM that has a propensity to form coiled-coils (Kawasaki, Lange, & Feske, 2009; Li et al., 2007; Muik et al., 2008; Park et al., 2009; Yang, Jin, Cai, Li, & Shen, 2012). Mutation of the residues corresponding to Ile316 and Leu319 of human Orai1, which mediate the coiled-coil packing in the crystal structures, prevents the interaction of Orai1 with STIM1 and subsequent channel activation (L273S and L276D mutations of human Orai1) (Muik et al., 2008; Navarro-Borelly et al., 2008). In accord with the unlatching we observe, an additional body of evidence suggests that the M4-ext helices undergo conformational changes that lead to channel activation (Navarro-Borelly et al., 2008; Palty, Fu, & Isacoff, 2017; Tirado-Lee, Yamashita, & Prakriya, 2015; Zhou et al., 2016). One possible mechanism of STIM binding is that unlatching would expose the M4-ext regions and make them available for interaction with STIM. Our observation that unlatching does not necessarily open the pore is consistent with studies indicating that STIM1 can bind to loss-of-function mutants of human Orai1 that have constitutively closed pores, such as the pore-lining R91W mutant (Derler et al., 2009; B. A. McNally, Somasundaram, Jairaman, Yamashita, & Prakriya, 2013). Congruently, we observe that the corresponding K163W mutant of *Drosophila* Orai can adopt an unlatched conformation without opening of the pore.

While the most pronounced structural differences between the closed and open pores are within the basic region, we find that removal of the basic region by mutation does not form a constitutively open channel. On the other hand, mutations within the hydrophobic region of the pore (e.g. F99C or V102A of human Orai1 or V174A of Orai) give rise to leaky channels, albeit with diminished selectivity for Ca^2+^ (Figure 8) (Hou et al., 2012; Beth A McNally et al., 2012; Yamashita et al., 2017). We hypothesize that the modest widening of the hydrophobic region observed in the open conformation, which accompanies the dramatic dilation of the basic region, is critical for ion permeation and that the hydrophobic region functions as a “gate” – a variable constriction that prevents or permits ion conduction. This hypothesis is consistent with the observation that hydrophobic substitutions of the upper basic residue (R91W in human Orai1 or K163W in Orai) create constitutively closed channels because hydrophobic packing in the basic region would likely prevent the dilation of the opened pore. Both structural and functional analyses point to the conclusion that while the conformational changes in the pore are more dramatic on the cytosolic side, they also extend to the extracellular side.

Ion binding in the open pore suggests that direct coordination of Ca^2+^ by the ring of Glu178 residues in the selectivity filter is responsible for the channel’s exquisite selectivity for Ca^2+^. The presence of a short selectivity filter, in this case just the single ring of glutamate residues, differentiates Orai from most other cation channels. Cation-selective channels that share a general architecture that was first identified by structure of the tetrameric potassium channel KcsA, which include voltage-dependent K^+^ channels, voltage-dependent Ca^2+^ channels, the Ca^2+^-selective channel TRPV6, and voltage-dependent Na^+^ channels, have longer selectivity filter regions that typically involve multiple ion-binding sites arranged in single file (Doyle et al., 1998; Liao, Cao, Julius, & Cheng, 2013; Morais-Cabral, Zhou, & Mackinnon, 2001; Payandeh, Scheuer, Zheng, & Catterall, 2011; Saotome, Singh, Yelshanskaya, & Sobolevsky, 2016; Tang et al., 2014; Wu et al., 2016). The presence of multiple ions in single file can provide ion-selectivity in the context of high conductivity of ions through the selectivity filter (Hille, 1992). Orai conducts Ca^2+^ very slowly in comparison to these channels, and its short selectivity filter may be one reason. In the physiological context of approximately 2 mM Ca^2+^ outside the cell, we hypothesize an ion-selectivity mechanism in which the selectivity filter toggles between having one and having two Ca^2+^ ions present. When one Ca^2+^ ion is bound, it would likely be centered within the glutamate ring. When two Ca^2+^ ions are present, the flexibility afforded by glutamate side chains may allow the ions to be positioned in single-file within the selectivity filter with one above the other. In this metastable state, the ion-ion repulsion would be sufficient to allow the lower Ca^2+^ ion to dissociate from the filter and move through the pore. The single Ca^2+^ ion remaining in the filter would not be easily displaced without the binding of a second Ca^2+^. The single Ca^2+^ ion, however, would block monovalent cations from permeating through the filter. On the other hand, when Ca^2+^ is artificially stripped away using a chelator, monovalent cations could stream through the selectivity filter unimpeded. The proposed mechanism would explain why micromolar concentrations of Ca^2+^ block monovalent cations from permeating through the pore (Hoth & Penner, 1993; Lepple-Wienhues & Cahalan, 1996). And it would explain why millimolar concentrations of Ca^2+^ are needed for efficient Ca^2+^ conduction because a high concentration of Ca^2+^ would favor transient binding of a second Ca^2+^ ion. Unlike the selectivity filters of most other cation channels that are designed for high ion-throughput, the selectivity filter of Orai may impose an energy barrier that impedes Ca^2+^ permeation. Other energy barriers that might serve to limit the flow of Ca^2+^ through the pore so as to not overwhelm the cell with Ca^2+^ include the narrow hydrophobic region and possible electrostatic repulsion by the basic region.

Comparison of the structures engenders a sequence for channel activation that proceeds from a quiescent state prior to interaction with STIM, through an unlatched intermediate, and culminates with an open pore (Figure 10, Movie 2). In the quiescent conformation, clasped latches constrain the outer cytosolic diameter of the channel and hold the pore closed. Unlatching, which could happen transiently and spontaneously, would expose cytosolic docking sites for STIM. The engagement of STIM, via molecular interactions that remain to be resolved, stabilizes an unlatched conformation and the widening of the pore that permits Ca^2+^ influx. The structures give insight into the remarkable molecular choreography by which Orai governs store-operated Ca^2+^ entry and a myriad of downstream cellular responses.

**Figure 10.**
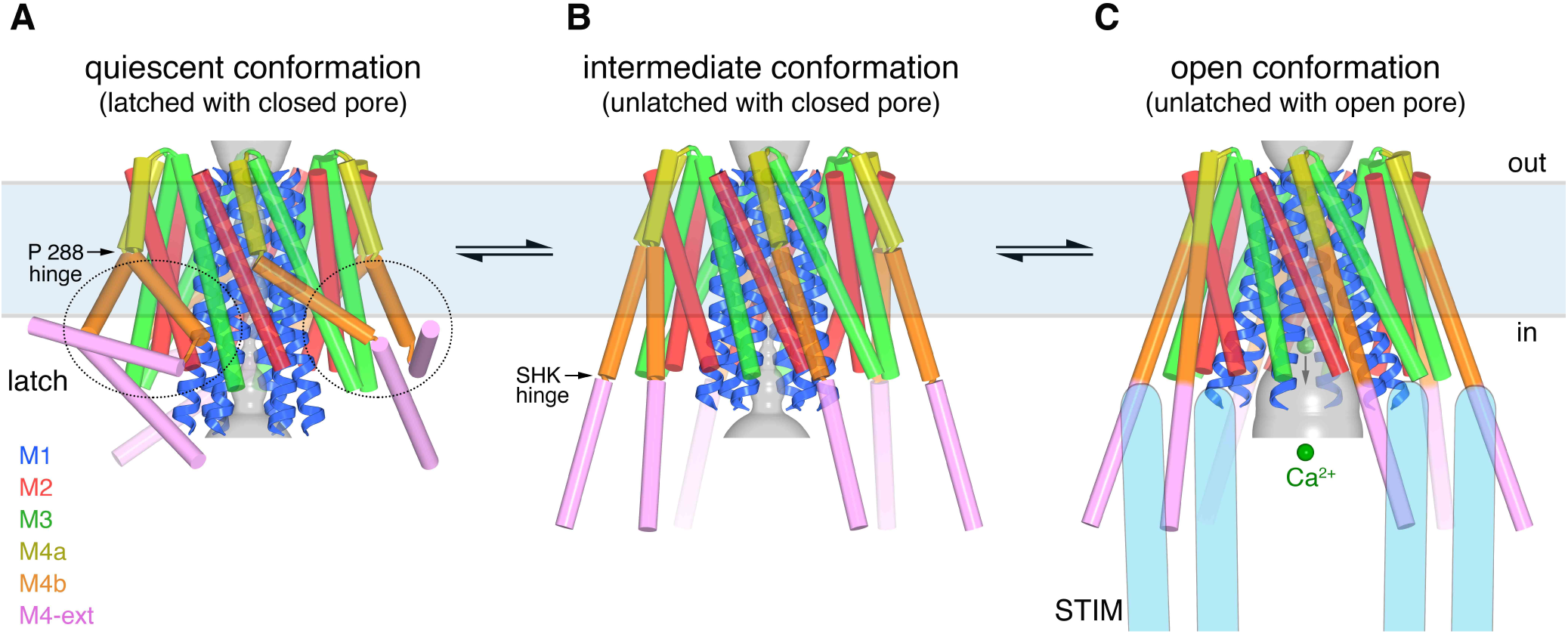
Proposed sequence of channel activation. **a**, Quiescent conformation of Orai prior to binding of STIM (from PDB ID 4HKR, cylinders and ribbons). The pore (gray surface) is closed and the latches are fastened (two latches are indicated with dashed ovals). The M4 helices are bent at Pro288, delineating them into M4a and M4b. The M4b portions (orange) interact with the M3 helices (green), in six-fold fashion, and prevent the pore from opening by constraining the cytosolic region of the M3 helices. The interaction between M4b and M3 and the bend at Pro288 are stabilized by three sets of paired M4-ext helices. **b**, An intermediate conformation: structure of WT Orai_cryst_ in which the pore is closed but the latches are released. Conformations of M1-M4a are indistinguishable from (**a**) (Figure 8D). When unlatched, mobile M4-ext regions are hypothesized to be available to interact with cytosolic regions of STIM that would become exposed as a result of depletion of Ca^2+^ from the ER. Spontaneous unlatching would not necessarily require STIM binding and does not necessarily open the pore. **c**, Open conformation. The structure of H206A Orai_cryst_ is shown (cylinders and ribbons), with approximate dimensions of the pore shown as a gray surface. Following store depletion, we hypothesize that STIM (blue shapes) engages with cytosolic regions of Orai and stabilizes the pore in an open conformation. On the basis of the effects of the H206A mutation, we suspect that the free energy difference between the intermediate and open conformations is on the order of only a few hydrogen bonds. Unlatching is required to allow the widening of the pore and the influx of Ca^2+^ (green spheres). Arrows between conformations denote equilibria and the horizontal rectangle indicates approximate boundaries of the plasma membrane. The depiction of the cytosolic region of STIM is conceptual and is not meant to imply stoichiometry or conformation. **Movie 1.** Electron density for the open conformation from Figure 3. **Movie 2.** Video showing opening sequence as illustrated in Figure 10.

## Materials and Methods

### Cloning, expression, purification and crystallization

cDNA encoding *Drosophila melanogaster* Orai (amino acids 133-341) followed by a C-terminal YL½ antibody affinity tag (amino acids EGEEF)(Kilmartin, Wright, & Milstein, 1982) was cloned into the EcoRI and NotI restriction sites of the *Pichia pastoris* expression vector pPICZ-C (Invitrogen Life Technologies). Two non-conserved cysteine residues were mutated to improve protein stability (C224S and C283T). This construct, termed “WT Orai_cryst_”, differs from the one we used previously (Hou et al., 2012) only in that it contains wild type Pro276 and Pro277 residues in the hyper variable M3-M4 loop rather than arginine substitutions at these positions. Constructs bearing the H206A or K163W mutations were made on the background of WT Orai_cryst_ using standard molecular biology techniques (designated “H206A Orai_cryst_” or “K163W Orai_cryst_”, accordingly). Transformations into *P. pastoris*, expression, and cell lysis were performed as previously described (Long, Campbell, & MacKinnon, 2005).

Lysed *P. pastoris* cells were re-suspended in buffer (3.3 ml buffer for each 1 g of cells) containing 150 mM KCl, 10 mM sodium phosphate, pH 7.0, 0.1 mg/ml deoxyribonuclease I (Sigma-Aldrich), 1:1000 dilution of Protease Inhibitor Cocktail Set III, EDTA free (CalBiochem), 1 mM benzamadine (Sigma-Aldrich), 0.5 mM 4-(2-aminoethyl) benzenesulfonyl fluoride hydrochloride (Gold Biotechnology) and 0.1 mg/ml soybean trypsin inhibitor (Sigma-Aldrich). Cell lysate was adjusted to pH 7.0 with 1 N KOH, 0.11 g n-dodecyl-β-D-maltopyranoside (DDM, Anatrace, solgrade) per 1 g of cells was added to the cell lysate, and the mixture was stirred at room temperature for 45 minutes to extract Orai from the membranes. The sample was then centrifuged at 30,000 *g* for 45 min at 17°C and the supernatant was filtered (0.45 μm polyethersulfone membrane). YL½ antibody (IgG, expressed from hybridoma cells and purified by ion exchange chromatography) was coupled to CNBr-activated sepharose beads (GE Healthcare) according to the manufacturer’s protocol. Approximately 0.4 ml of beads were added to the sample for each 1 g of *P. pastoris* cells and the mixture was rotated at room temperature for 1 h. Beads were collected on a column, washed with 5 column-volumes of buffer containing 150 mM KCl, 10 mM sodium phosphate, pH 7.0, 5 mM DDM, 0.1 mg/ml lipids (3:1:1 molar ratio of 1-palmitoyl-2-oleoyl-sn-glycero-3-phosphocholine, 1-Palmitoyl-2-oleoyl-sn-glycero-3-phosphoethanolamine, and 1-palmitoyl-2-oleoyl-sn-glycero-3-[phospho-rac-(1-glcerol)], obtained from Avanti) and eluted with buffer containing 150 mM KCl, 100 mM Tris-HCl, pH 8.5, 5 mM DDM, 0.1 mg/ml lipids and 5 mM Asp-Phe peptide (Sigma-Aldrich). The eluted protein was concentrated to ∼ 25 mg/ml using a 100 kDa concentrator (Amicon Ultra, Millipore) and further purified on a Superdex-200 gel filtration column (GE Healthcare) in 75 mM KCl, 10 mM Tris-HCl, pH 8.5, 0.1 mg/ml lipids, and detergent: 4 mM octyl glucose neopentyl glycol (Anatrace, anagrade) to obtain crystals of K163W Orai_cryst_ in space group P4_2_2_1_2, or a mixture of 0.5 mM decyl maltose neopentyl glycol (Anatrace, anagrade) and 3 mM octyl glucose neopentyl glycol for crystals of WT and K163W Orai_cryst_ in space group of I4_1_, and 0.5 mM decyl maltose neopentyl glycol for H206A Orai_cryst_. For crystals of H206A Orai_cryst_, 3 mM octyl glucose neopentyl glycol was added into the purified H206A Orai_cryst_ just before setting up crystallization trials. A typical prep, utilizing 20 g of cells, yielded ∼ 2 mg of purified Orai. For the crystals of H206A Orai_cryst_ with I^−^, NaI was substituted in place of KCl in the purification buffers. For crystals of H206A in Ba^2+^, 5 mM BaCl_2_ was added to the final purified protein before crystallization. Purified Orai proteins were concentrated to 10-20 mg/ml using 100 kDa Vivaspin-2 concentrators (Sartorius Stedim Biotech), and mixed 1:1 (250 nl: 250 nl) with crystallization solutions for hanging drop vapor diffusion crystallization. Crystals of WT Orai_cryst_ grew in 32-35% PEG 400 (v/v) and 0.1 M potassium phosphate pH 7.5. Crystals of K163W Orai_cryst_ in space group P4_2_2_1_2 grew in 24-26% PEG 400 (v/v) and 0.2 M potassium phosphate pH 6.5. Crystals of K163W Orai_cryst_ in space group I4_1_ grew in 26-28% PEG 400 (v/v) and 150 mM NaCl and 100 mM N-2-hydroxyethylpiperazine-N-2’-ethanesulfonic acid (HEPES) pH 7.5. Native crystals of H206A Orai_cryst_ grew in 36-38% PEG400, 500 mM NaCl and 100 mM Tris-HCl pH 9.0. The crystallization solution for H206A Orai_cryst_ with NaI was 28-31% PEG400 and 100 mM Tris-HCl pH7.5. The crystallization solution for H206A Orai_cryst_ with BaCl_2_ was 32-34% PEG400, 500 mM NaCl and 100 mM Tris-HCl pH 8.5.

## Structure determination

### Data collection and processing

All crystals were dehydrated and cryo-protected before flash-cooling in liquid nitrogen by serial transfer into solutions containing the buffer components of an equilibrated crystallization drop and increasing concentrations of PEG 400 (to 50% w/v) in 9 steps with ∼ 1 min intervals. For heavy atom derivatives, Crystals of K163W Orai_cryst_ belonging to space group I4_1_ were soaked in stabilization solution supplemented with ∼18 μg/ml p-chloromercuribenzene sulfate (PCMB), or ∼7 μg/ml di-μ-iodo-bis(ethylene-diamine)-di-platinum(II) nitrate (PIP) for 24 hours. For ion binding experiments, crystals of H206A Orai_cryst_ were soaked in stabilization solution supplemented with 1 mM GdCl_3_ for two days. After soaking, the crystals were cryo-protected in the same solutions as native crystals and flash-cooled. Crystals of H206A Orai_cryst_ that contained 5 mM BaCl_2_ were soaked in stabilization solution supplemented with 50 mM BaCl_2_ during dehydration steps and flash-cooled. X-ray diffraction data sets were collected using synchrotron radiation and were indexed, integrated and scaled with the HKL suite (Otwinowski & Minor, 1997) or XDS (Kabsch, 2010). Resolution limits of the diffraction data were estimated from the CC_1/2_ value (Karplus & Diederichs, 2012).

### K163W Orai_cryst_ (P4_2_2_1_2 space group)

Initial phases for data collected from crystals of K163W Orai_cryst_ belonging to space group P4_2_2_1_2, were determined by molecular replacement (MR) with PHENIX(Adams et al., 2010) using residues 148-288 of the structure of K163W *Drosophila* Orai in the quiescent conformation (PDB ID: 4HKS) as a search model. The asymmetric unit contains three Orai subunits; these form a complete hexameric channel by a two-fold rotational symmetry operator of the P4_2_2_1_2 space group. To improve the phases and reduce bias, the phases were improved with solvent flattening, histogram matching, and 3-fold non-crystallographic symmetry (NCS) averaging with the program DM(Cowtan, 1994). This yielded well-defined density for the channel (Figure 8-figure supplement 1). A B-factor sharpening value of −150 Å^2^ was applied to the electron density maps that are displayed (Figure 3, Figure 8, and Figure 8-figure supplement 1). The atomic model was adjusted in COOT (Emsley, Lohkamp, Scott, & Cowtan, 2010) and refined in CNS using a deformable elastic network (DEN) force field (Brünger et al., 1998; Brunger et al., 2012; Schröder, Brunger, & Levitt, 2007; Schröder, Levitt, & Brunger, 2010) and in PHENIX with NCS and secondary structure restraints. The final model contains residues 148-327 of Orai, excluding the following residues that did not have well-enough defined electron density to direct model building: 181-188 (the M1-M2 loop), 220-239 (the M2-M3 loop), and 314-327 of subunit B.

### K163W Orai_cryst_ (I4_1_ space group)

Initial phases for K163W Orai_cryst_ in space group I4_1_ were determined experimentally by the MIRAS (multiple isomorphous replacement with anomalous scattering) method using a native dataset and two derivative ones (PCMB and PIP) using SHARP(Vonrhein, Blanc, Roversi, & Bricogne, 2007) (Table 2; Figure 8-figure supplement 1E)). The asymmetric unit contains four hexameric channels (twenty four Orai subunits) for which density was apparent following solvent flattening with the program DM (Cowtan, 1994). The phases were improved and extended to 6.1 Å resolution using solvent flattening, histogram matching, and 24-fold NCS averaging with DM. This yielded continuous densities for all helices in all 24 Orai subunits within the asymmetric unit (Figure 8-figure supplement 1A). Anomalous-difference electron density maps of well-ordered platinum and mercury sites at Met321 and Cys215, respectively, helped establish the amino acid register. The quiescent conformation structure (PDB ID: 4HKS; determined at 3.35 Å) was used as a reference for modeling of the M1-M3 portion and the 4.35 Å structure in space group P4_2_2_1_2 was used for M4/M4-ext. Minor adjustments of the M4 and M4-ext helices were made as necessary. Refinement was done using rigid body and DEN refinement in CNS utilizing NCS, helical secondary structure (backbone phi, psi) and phase restrains(Brünger et al., 1998; Brunger et al., 2012; Schröder et al., 2007; Schröder et al., 2010). Grouped B-factor and TLS refinement were performed (in PHENIX), for which each of the four channels was defined as a group, as is appropriate for modest-resolution data. The final model contains residues 148-327 of Orai, excluding the following residues that did not have well-enough defined electron density to direct model building: 181-188 (the M1-M2 loop) and 220-239 (the M2-M3 loop).

### WT Orai_cryst_ (I4_1_ space group)

Initial phases for the structure of WT Orai_cryst_ were determined by MR using the K163W Orai_cryst_ structure (space group I4_1_) as an initial model in the program PHENIX (Adams et al., 2010). Diffraction data were collected to maximize the anomalous scattering from iron (λ = 1.738 Å). Single-wavelength anomalous diffraction (SAD) phases derived from the anomalous density (presumably from iron) in the basic region of the pore were combined with the molecular replacement phases (MR-SAD method) using AutoSol of PHENIX (Adams et al., 2010). Potential phase bias was further minimized by density modification with solvent flattening, histogram matching, and 24-fold NCS averaging using the program DM (Cowtan, 1994). Model building was aided by the quiescent conformation (PDB ID: 4HKR; 3.35 Å resolution) and by the 4.35 Å resolution structure of K163W Orai_cryst_ from the P4_2_2_1_2 space group. Minor adjustments of the M4-ext helices were made in COOT (Emsley et al., 2010). Refinement was done using rigid body and DEN refinement in CNS (Brünger et al., 1998; Brunger et al., 2012; Schröder et al., 2007; Schröder et al., 2010). During refinement, NCS, helical secondary structure (backbone phi, psi) and experimental phase restrains were applied. Grouped B-factor and TLS refinement were performed (in PHENIX), for which each of the four channels was defined as a group. The final model contains residues 148-327 of Orai, excluding the following residues that did not have well-enough defined electron density to direct model building: 181-191 (the M1-M2 loop) and 220-239 (the M2-M3 loop).

### H206A Orai_cryst_ (I4_1_ space group)

Initial phases for the structure of H206A Orai_cryst_ were obtained with MR using a truncated structure (amino acids 148-309) of WT Orai_cryst_ (I4_1_ space group) as a search model in PHENIX. At this stage electron density maps contained broken density for the four channels in the asymmetric unit. These phases were used to identify four Gd^3+^ sites (one site in the glutamate ring of each channel in the ASU) from the dataset collected from a crystal soaked in GdCl_3_ (Table 1) and the phases were improved using the MR-SAD method (using AutoSol in PHENIX). The phases were then improved using solvent flattening, histogram matching, and four-fold non-crystallographic symmetry (NCS) averaging with the program DM(Cowtan, 1994) using an entire channel as the reference region for NCS averaging. This yielded continuous electron density for helices. These phases were used as starting phases for the native dataset and were improved and extended to 6.7 Å resolution using the 24-fold NCS present within the asymmetric unit. For the 24-fold averaging, a single Orai subunit corresponding to amino acids 148-309 was used as the reference region (with solvent flattening, histogram matching, and NCS averaging performed in DM). This map (shown in Figure 3A,B) was used to direct model building. The initial model was generated by rigid body fit of WT Orai_cryst_ subunits and adjusted manually in COOT (e.g. to account for the additional bend in M1)(Emsley et al., 2010). Side chain conformations cannot be determined from the electron density due to the limit of the diffraction data; side chains are included in the atomic model for reference, however, and their conformations are based on those observed in the quiescent conformation (PDB 4HKR) for amino acids in M1-M4a and those from the 4.35 Å resolution structure presented here (P4_2_2_1_2 space group of K163W Orai_cryst_) for amino acids in M4b and M4-ext. Helical regions were modeled with ideal α-helical geometry and side chain rotamers were selected from frequently occurring conformations (Hintze, Lewis, Richardson, & Richardson, 2016) and to minimize steric clashes (Word et al., 1999). Refinement was done using rigid body and DEN refinement in CNS utilizing NCS and helical secondary structure (backbone phi, psi) restrains (Brünger et al., 1998; Brunger et al., 2012; Schröder et al., 2007; Schröder et al., 2010). Grouped B-factor and TLS refinement were performed (in PHENIX), for which each of the four channels was defined as a group. Highly redundant data allowed us to visualize anomalous-difference electron density arising from the sulfur atoms of methionine or cysteine residues on each of the M1-M4 helices and on the M4-ext helix (Table 1). These anomalous peaks confirm the assigned amino acid register and indicate the accuracy of the crystallographic phases (Figure 3-figure supplement 1). The model contains residues 148-327 of Orai, excluding the following residues that did not have well-enough defined electron density to direct model building: 181-191 (the M1-M2 loop) and 220-239 (the M2-M3 loop).

### Anomalous-*difference* electron density maps for ion experiments

Phases for the three anomalous-difference electron density maps in Figure 5 (Table 1) were determined by MR-SAD (Phenix, using H206A Orai_cryst_ for MR and the anomalous signal from ions), and were improved by 24-fold NCS averaging, solvent flattening and histogram matching in DM. Anomalous difference electron density for each ion was observed, with approximately the same sigma level and position, in all four channels of each asymmetric unit.

#### Reconstitution and flux assay

Orai constructs were purified and reconstituted into lipid vesicles using a modified published procedure(Hou et al., 2012). A lipid mixture containing 15 mg/ml POPE (1-palmitoyl-2-oleoyl-sn-glycero-3-phosphocholine) and 5 mg/ml POPG (1-palmitoyl-2-oleoyl-sn-glycero-3-phospho(1’-rac-glycerol)) was prepared in water and solubilized with 8% (w/vol) n-decyl-β-D-maltopyranoside. Purified WT or H206A Orai_cryst_ protein was mixed with the solubilized lipids to obtain a final protein concentration of 0.5 mg/ml and a lipid concentration of 10 mg/ml. Detergent was removed by dialysis (15 kDa molecular weight cutoff) at 4 °C for 7 days against a reconstitution buffer containing 10 mM HEPES pH 7.0, 150 mM KCl and 0.2 mM ethylene glycol tetraacetic acid (EGTA), with daily buffer exchanges and utilizing a total volume of 14 l of reconstitution buffer. The reconstituted sample was sonicated (∼30 sec), aliquoted, flash-frozen in liquid nitrogen and stored at −80 °C.

Vesicles were rapidly thawed (using 37 °C water bath), sonicated for 5 sec, incubated at room temperature for 2-4 hours before use, and then diluted 100-fold into a flux assay buffer containing 150 mM n-methyl-d-glucamine (NMDG), 10 mM HEPES pH 7.0, 0.2 mM EGTA, 0.5 mg/mL bovine serum albumin and 0.2 μM 9-amino-6-chloro-2-methoxyacridine (ACMA, from a 2 mM stock in DMSO). Data were collected on a SpectraMax M5 fluorometer (Molecular Devices) using Softmax Pro 5 software. Fluorescence intensity measurements were collected every 30 sec over the span of the 1200 sec experiment (excitation and emission set to 410 nm and 490 nm, respectively). The proton ionophore carbonyl cyanide m-chlorophenyl hydrazine (CCCP, 1 μM from a 1 mM stock in DMSO) was added after 150 sec and the sample was mixed briefly by pipette. The potassium ionophore valinomycin (2 nM from a 2 uM stock in DMSO) was added at the end of the experiment (990 sec) to establish a baseline fluorescence and confirm vesicle integrity. For experiments used to determine the effects of Ca^2+^, Mg^2+^ or Gd^3+^ on H206A Orai_cryst_, CaCl_2_ (1 mM final concentration), MgCl_2_ (3 mM final concentration), or GdCl_3_ (0.1 mM final concentration) was added to aliquots of reconstituted vesicles. To introduce the ions into the vesicles, these vesicles were sonicated for 5 sec, frozen in liquid nitrogen, thawed, sonicated a second time for 5 sec, and incubated at room temperature for 2-4 hours prior to measurements. The flux assay buffers were supplemented with 1 mM CaCl_2_, 3 mM MgCl_2_, or 0.1 mM GdCl_3_, respectively, for these experiments.

Flux experiments using V174A Orai_cryst_ and Orai bearing the simultaneous R155S, K159S, and K163S mutations (SSS Orai_cryst_) were performed analogously. In these experiments Na^+^ flux was measured under divalent-free conditions as described (Hou et al., 2012). Purified protein was prepared as described above. Proteoliposomes, or liposomes without protein, were formed by dialysis against 10 mM HEPES, pH 7.0, 150 mM NaCl and 0.2 mM EGTA, and were aliquoted, flash-frozen in liquid nitrogen and stored at −80 °C until use. For reconstitution, the lipid concentration was 10 mg/ml POPE:POPG (3:1 weight ratio) and the protein concentrations were 1 mg/ml and 0.01 mg/ml for SSS Orai_cryst_ and V174A Orai_cryst_, respectively. (0.1 and 0.01 mg/ml SSS Orai_cryst_ concentrations were also tested and gave indistinguishable results.) Liposome samples were diluted 50-fold into flux buffer containing 10 mM HEPES pH 7.0, 0.2 mM EGTA, 0.5 mg/mL bovine serum albumin, 0.2 μM ACMA, and 150 mM N-methyl-D-glucamine (NMDG), which established a Na^+^ gradient. After stabilization of the fluorescence signal (150 sec), 1 μM CCCP was added to the sample. The Na^+^ ionophore monensin was added after 990 sec to render all vesicles permeable to Na^+^ and establish the minimum baseline fluorescence.

## Acknowledgments

We thank Richard Hite, Christopher Lima, Nikola Pavletich, and members of the Long laboratory for discussions and comments on the manuscript. This work was supported by NIH Grant R01 GM094273 (to S.B.L.) and a core facilities support grant to Memorial Sloan Kettering Cancer Center (P30 CA008748). Synchrotron radiation facilities were supported by NIH Grants ACB-12002, AGM-12006, P41 GM103403, and S10 RR029205, under Department of Energy Contract DE-AC02-06CH11357. Atomic coordinates, structure factors, and crystallographic phases have been deposited in the Protein Data Bank with accession numbers 6BBF (H206A Orai_cryst_), 6BBG (WT Orai_cryst_), 6BBH (K163W Orai_cryst_, I4_1_ form), and 6BBI (K163W Orai_cryst_, P4_2_2_1_2 form).

## Author Contributions

X.H. and S.B. expressed, purified and crystallized proteins, and performed functional assays. All authors contributed to experimental design. X.H. determined structures. S.B.L. and X.H. prepared the manuscript.

---

**Figure 3-figure supplement 1.**
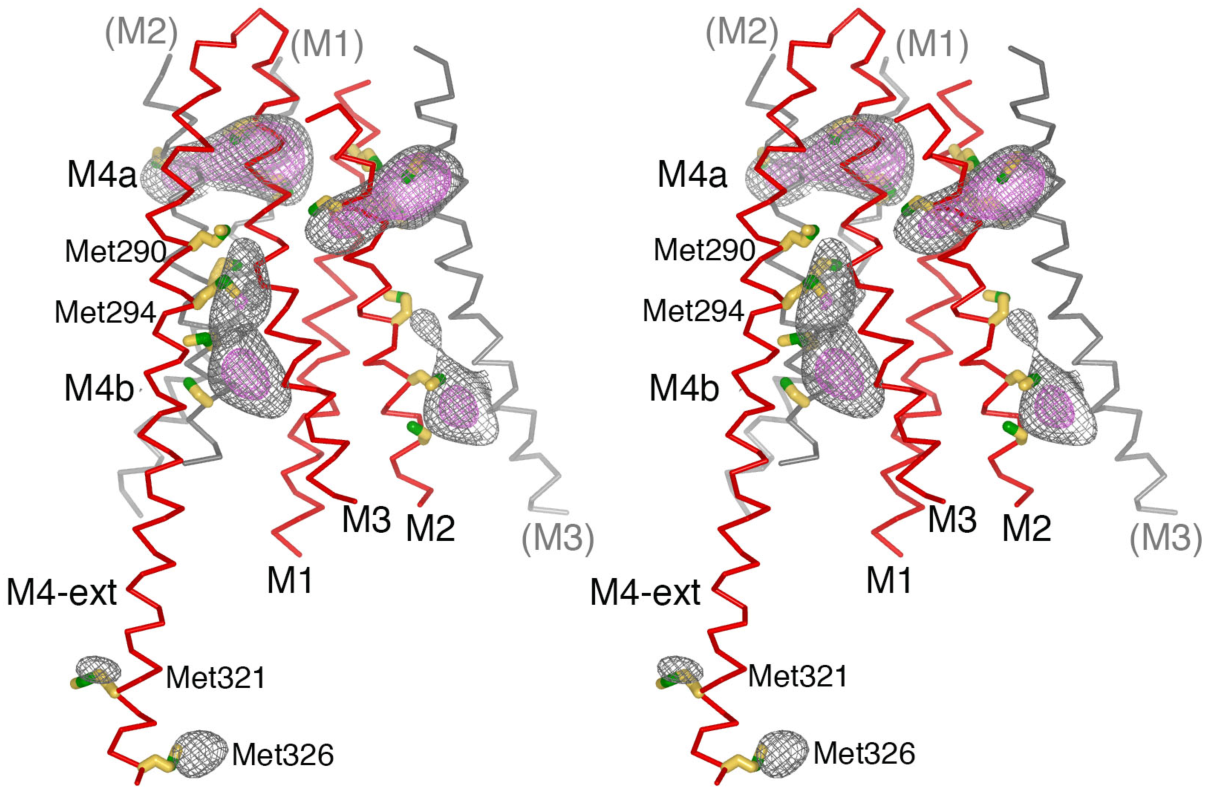
Anomalous-difference electron-density at cysteine and methionine residues in the final model of H206A Orai_cryst_, depicted in stereo. An anomalous-difference electron-density map was calculated from 25 to 10 Å resolution from highly redundant diffraction data collected with λ = 1.7085 Å X-rays (NaI experiment, Table 1) using anomalous differences as amplitudes and phases that were determined by MR-SAD, 24-fold NCS averaging, solvent flattening and histogram matching (Methods). This map was then averaged in real-space according to the 24-fold NCS symmetry to yield the map shown. The map is contoured at 5.5 σ (gray mesh) and 8.5 σ (pink mesh) and shown in the vicinity of a subunit of Orai (red Cα trace). Methionine and cysteine residues are shown as sticks (colored yellow for carbon and green for sulfur atoms). Methionine residues on M4b and M4-ext are labeled. Portions of neighboring Orai subunits (gray Cα traces) are shown for reference with their helices labeled in parentheses. While their side chain conformations are hypothetical on account of the limited resolution of the diffraction data, anomalous-difference electron-density peaks for methionine and/or cysteine residues on each of the M1-M4 helices and on the M4-ext helix confirm the amino acid register of the atomic model.

**Figure 4-figure supplement 1.**
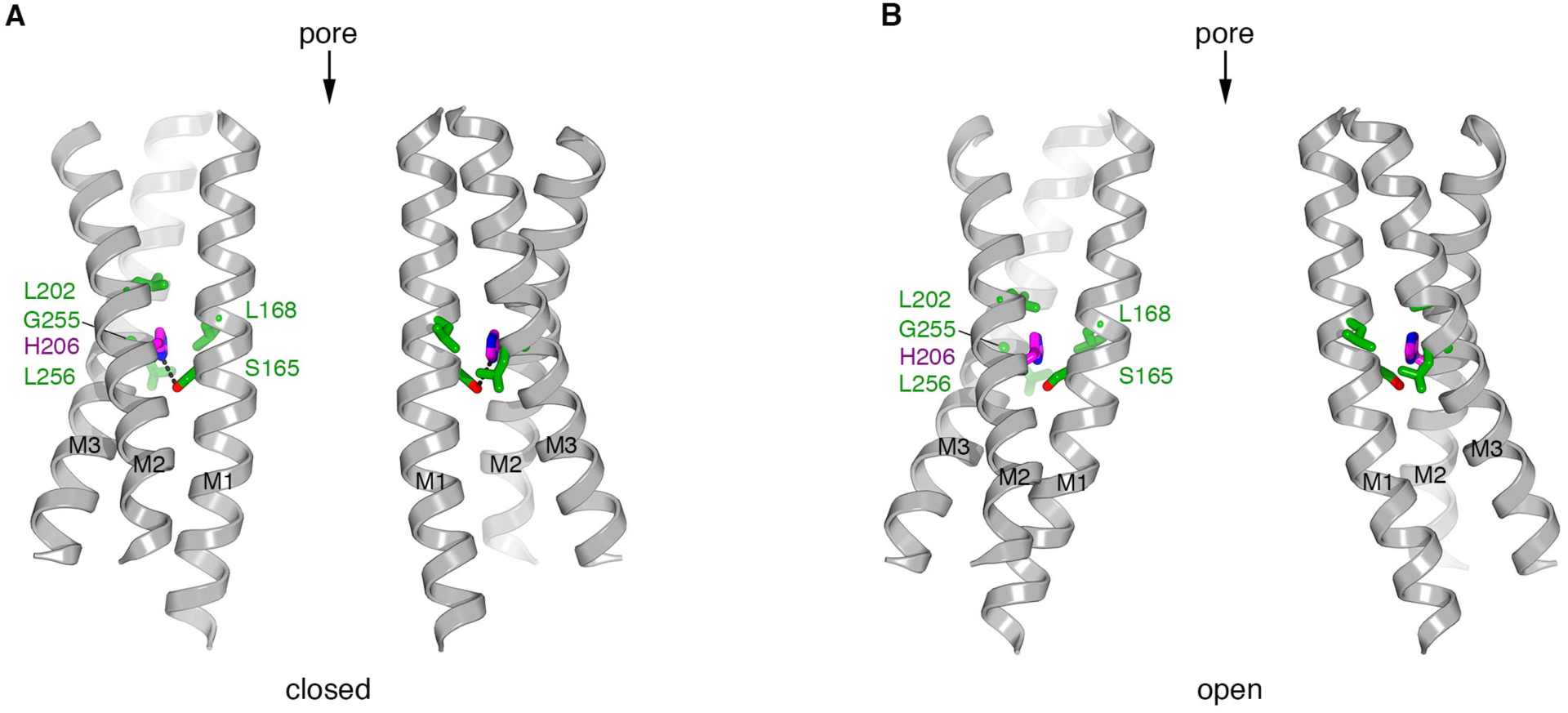
Residue 206 in closed and open pores. **a**, Depiction of the closed pore (PDB ID 4HKR). M1 through M3 are drawn as ribbons for two apposing subunits. His206 (H206, pink) and the amino acid side chains within van der Waals distance (green) are drawn as sticks. A hydrogen bond made between His206 and Ser165 is shown as a dashed line. **b**, Depiction of the open conformation (H206A Orai_cryst_), showing the corresponding regions as in (**a**). The conformations of the amino acid side chains in the atomic model are shown for reference to indicate plausibility despite the limited resolution of the diffraction data. Amino acid 206 is depicted as the wild type histidine to indicate that this amino acid could be accommodated in the observed conformation of the channel without steric hindrance.

**Figure 5-figure supplement 1.**
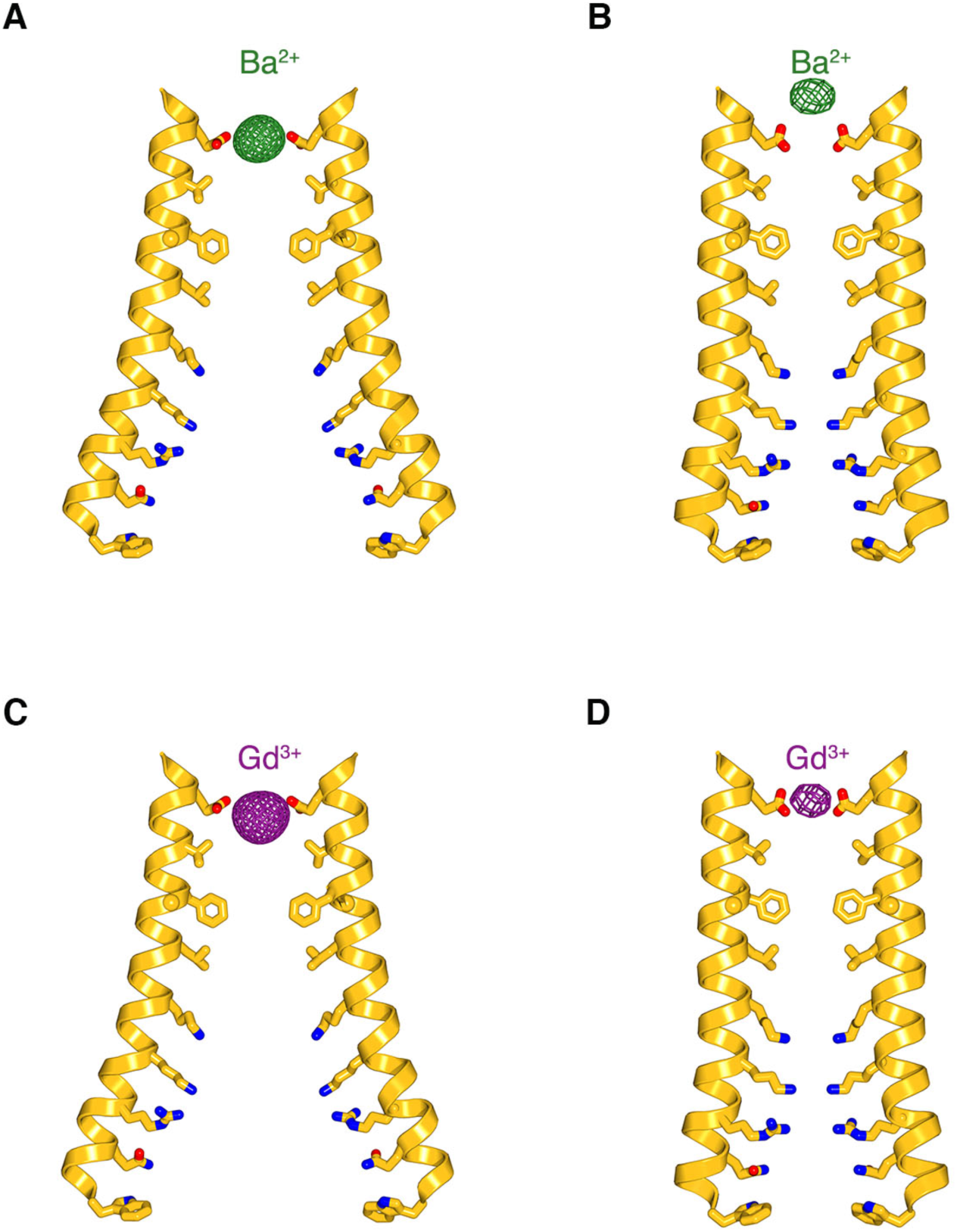
Comparison of the anomalous-difference electron-density peaks for Ba^2+^ and Gd^3+^ in the open and closed pores. **a-b**, Anomalous-difference electron density for Ba^2+^ (green mesh) in open pore of H206A Orai_cryst_ (from Figure 5A) and in the closed pore of the quiescent conformation (from (Hou et al., 2012)), respectively. M1 helices (amino acids 148 to 180) are depicted from two opposite subunits as ribbon representations. Pore-lining side chains (sticks) are drawn for reference in (**a**) since their conformations cannot be determined due to the limits of the diffraction data, and they are shown in (**b**) according to their observed conformations (Hou et al., 2012). **c-d**, Analogous depictions for the anomalous-difference electron-density peaks for Gd^3+^ in the open pore (**c**, from Figure 5B) and closed pore (**d**, from (Hou et al., 2012)).

**Figure 7-figure supplement 1.**
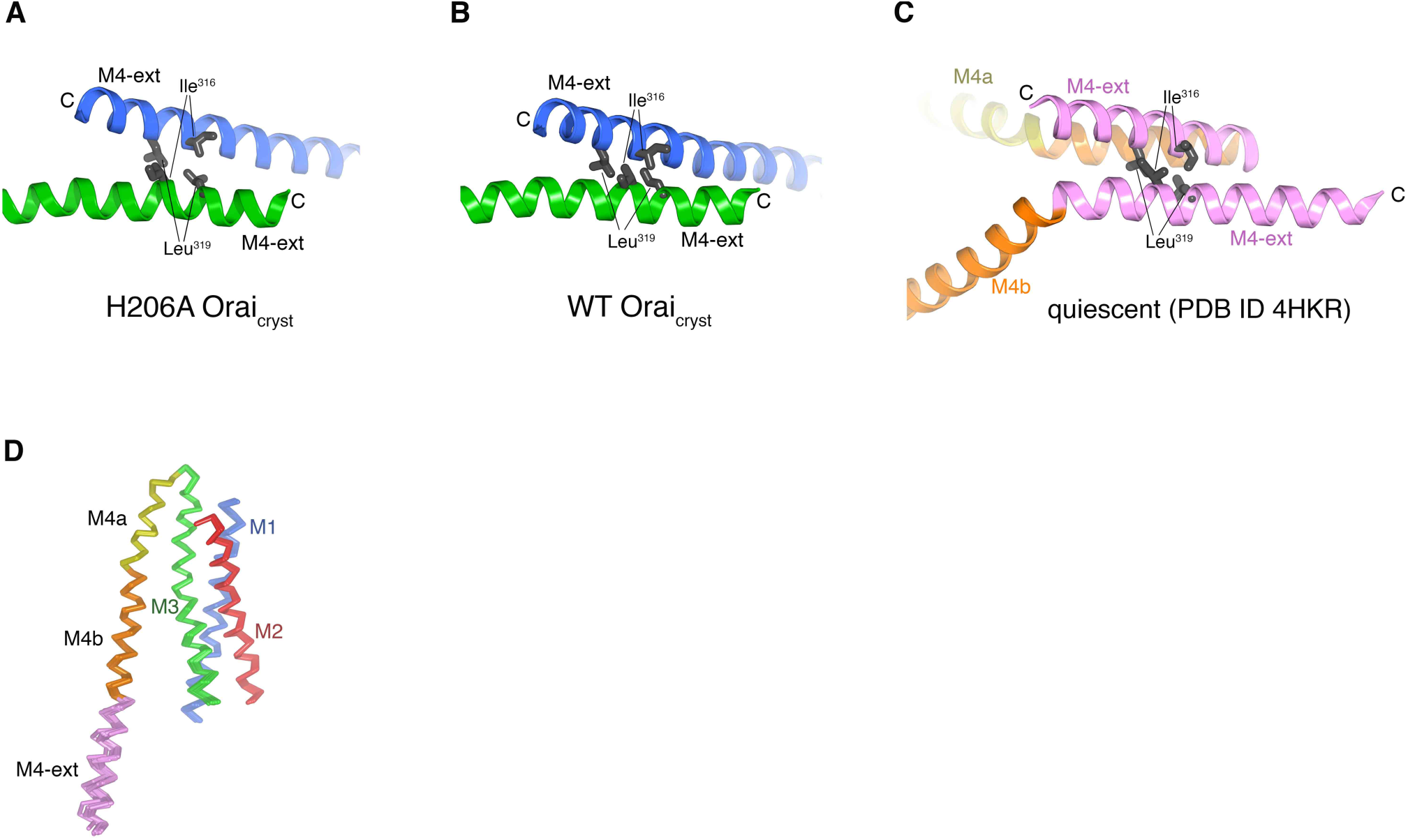
M4-ext helices and subunit comparisons. **a**, Close up view of a coiled-coil interaction between two M4-ext helices of the blue- and green-colored channels in the asymmetric unit of H206A Orai_cryst_ (from Figure 7A). Ile316 and Leu319, which form the hydrophobic interface of the coiled-coil interaction on each of the M4-ext helices, are drawn as gray sticks for reference. **b**, Analogous coiled-coil interaction between two M4-ext helices of the blue- and green- colored channels in the asymmetric unit of WT Orai_cryst_ (from Figure 7B). **c**, Coiled-coil interaction of two paired M4-ext helices observed in the quiescent conformation, which occurs between adjacent subunits of the same channel (from Figure 1, PDB ID 4HKR). **d**, Superposition of the 24 individual subunits (from four hexameric channels) of H206A Orai_cryst_ within the asymmetric unit, shown in Cα representation.

**Figure 8-figure supplement 1.**
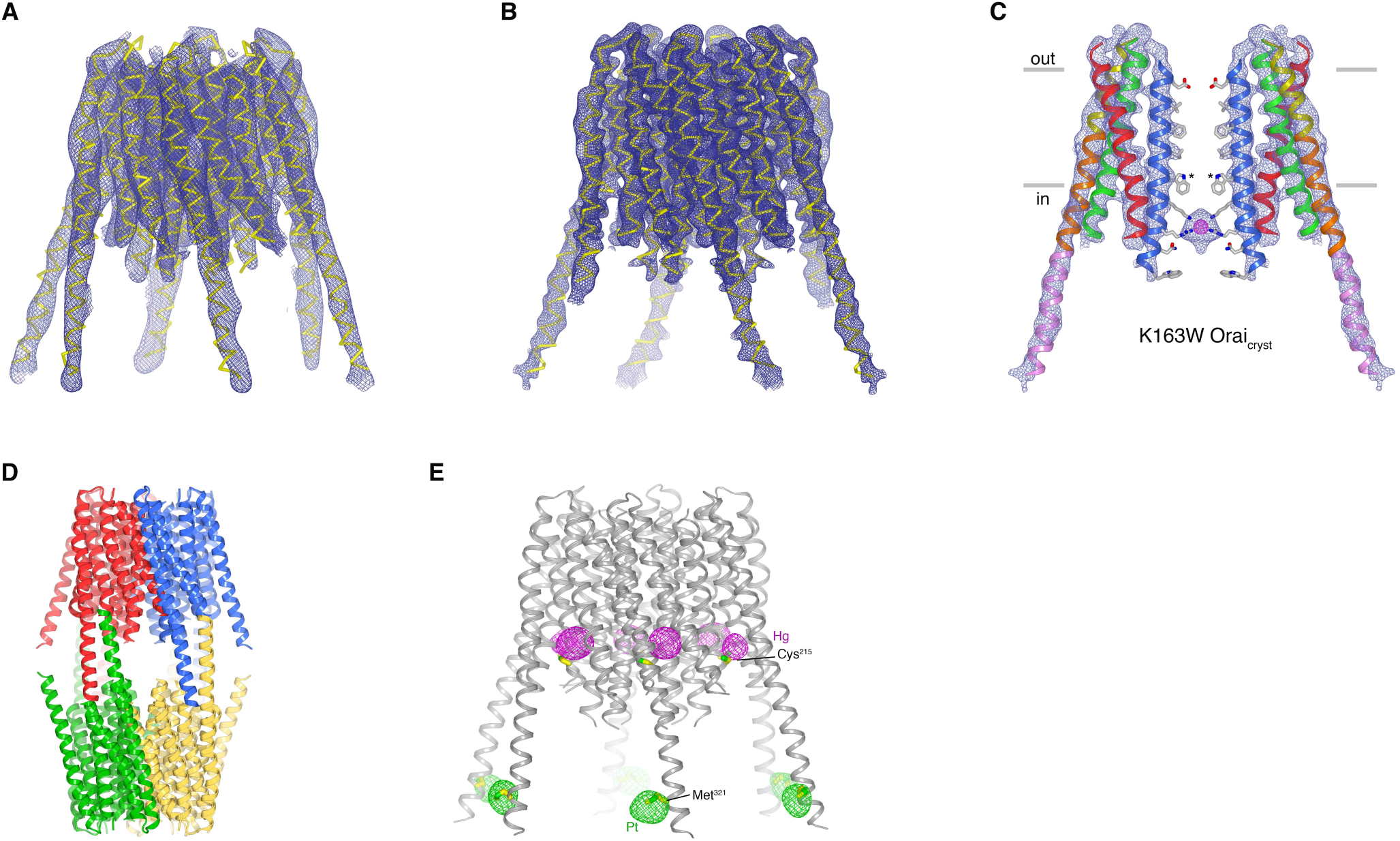
Structures of K163W Orai_cryst_ in I4_1_ and P4_2_2_1_2 crystal forms (unlatched with closed pore). **a**, Structure of K163W Orai_cryst_ from the I4_1_ crystal form. The map (contoured at 1.3 σ) was calculated from 20 – 6.1 Å using native-sharpened amplitudes and experimental phases that were determined by MIRAS and were improved by 24-fold NCS averaging, solvent flattening and histogram matching (Methods). The atomic model is shown in Cα representation (yellow). The crystal form is analogous to crystals of H206A Orai_cryst_ and WT Orai_cryst_, and has analogous crystal packing. The root-mean-squared deviation (RMSD) for Cα positions between the structures of WT Orai_cryst_ and K163W Orai_cryst_ is 0.5 Å. **b**, 4.35 Å resolution structure of K163W Orai_cryst_ in the P4_2_2_1_2 crystal form. Electron density (mesh) covering the channel is shown. The map (contoured at 1.3 σ) was calculated from 20 – 4.35 Å using sharpened amplitudes and phases that were determined by MR and were improved by 3-fold non-crystallographic symmetry (NCS) averaging, solvent flattening and histogram matching (Methods). **c**, Two opposing subunits of K163W Orai_cryst_ (P4_2_2_1_2 crystal form), showing the pore, with electron density from (**b**). Amino acids on the pore are depicted as sticks (conformations based on PDB 4HKS; Methods). Anomalous-difference electron density in the pore is shown as magenta mesh (calculated from 30 - 8 Å resolution using anomalous differences as amplitudes, and contoured at 3.8 σ). Asterisks mark the locations of K163W substitutions. **d**, Packing of K163W Orai_cryst_ in the P4_2_2_1_2 crystal form. The contents of each asymmetric unit (three Orai subunits) are colored a unique color. Two asymmetric units (e.g. blue and red) form a complete channel. The channels interact with one another in the crystal lattice via coiled-coil interactions between their M4-ext helices. **e**, Anomalous-difference electron-density for heavy atom derivatives of K163W Orai_cryst_ in the I4_1_ space group. One channel of the asymmetric unit is depicted as ribbons. The map for the platinum (Pt) derivative (magenta mesh, calculated from 25 to 8.0 Å, and contoured at 4.5 σ) was calculated from data collected from a crystal soaked in PIP (Table 2), using anomalous differences as amplitudes and phases from **a**. The analogous map for the mercury (Hg) derivative (green mesh, calculated from 25 to 8.0 Å, and contoured at 5 σ) was calculated from data collected from a crystal soaked in PCMB (Table 2). Each channel in the asymmetric unit has anomalous-difference density at these sites (24 sites for each derivative, Table 2). Cys^215^ and Met^321^ residues, to which the heavy atoms presumably bind, are depicted as sticks.

